# Development of Second-Generation Acyl Silane Photoaffinity Probes for Cellular Chemoproteomic Profiling

**DOI:** 10.1101/2025.05.24.655972

**Authors:** Annika C. S. Page, Lauren M. Orr, Margot L. Meyers, Bridget P. Belcher, Theodore G. Coffey, Spencer O. Scholz, Sabine Cismoski, Daniel K. Nomura, F. Dean Toste

## Abstract

Deconvolution of the protein targets of hit compounds from phenotypic screens, often conducted in live cells, is critical for understanding mechanism of action and identifying potentially hazardous off-target interactions. While photoaffinity labeling and chemoproteomics are long-established approaches for discovering small-molecule-protein interactions in live cells, there are a relatively small number of photoaffinity labeling strategies that can be applied in intracellular settings. Recently, we reported a novel chemical framework for photoaffinity labeling based on the photo-Brook rearrangement of acyl silanes and demonstrated its ability, when appended to protein-targeting ligands, to label recombinant proteins. Here, we report the application of these probes to live cell photoaffinity workflows, demonstrate their complementarity to current state-of-the-art minimalist diazirine-based photoaffinity probes, and introduce a modular synthetic route to access acyl silane scaffolds with improved labeling properties.

## Introduction

Chemoproteomics, particularly photoaffinity labeling (PAL), is a powerful approach for mapping small-molecule-protein interactions within the native cellular context^1–8^. Identifying the protein targets of bioactive small molecules, especially those emerging from phenotypic screens, is critical for elucidating mechanisms of action and anticipating potential off-target effects^1–8^. Current photoaffinity labeling methods predominantly employ reactive moieties such as benzophenones, aryl azides, and diazirines, each offering distinct advantages and limitations. Benzophenones, while highly stable and easily activated by UV light, often require prolonged irradiation and exhibit limited specificity due to nonspecific labeling. Aryl azides feature relatively simple synthesis and efficient crosslinking; however, their short-lived and highly reactive nitrene intermediates frequently lead to nonspecific protein labeling and lower yields. Diazirines, particularly minimalist diazirines, have gained popularity due to their compact size, efficient photolysis to highly reactive carbenes, and enhanced labeling selectivity^1–8^ In particular, linear aliphatic diazirines have been employed as so-called “minimalist” probes due to their small size and have experienced widespread use in the community. However, challenges inherent to the diazirine functionality remain. While minimalist probes have resulted in many successful small-molecule target identifications, much of the labeling events observed through diazirine irradiation occurs through a diazo isomer, whose long lifetime in solution can lead to the capture of non-specific proteins after eventual formation of the carbene resulting in false positive hits as well as an overrepresentation of membrane-related proteins through protonation of the diazo species and subsequent labeling by acidic residues on those proteins^7,9^ **(Figure 1A).** Some efforts have been made to reduce labeling from the diazo species^8,10^, though most PAL campaigns still use the original minimalist scaffold. PAL handles that avoid undesired reactivity and enable cellular target identification would thus expand the available toolkit for target identification campaigns^11^.

**Figure 1.**
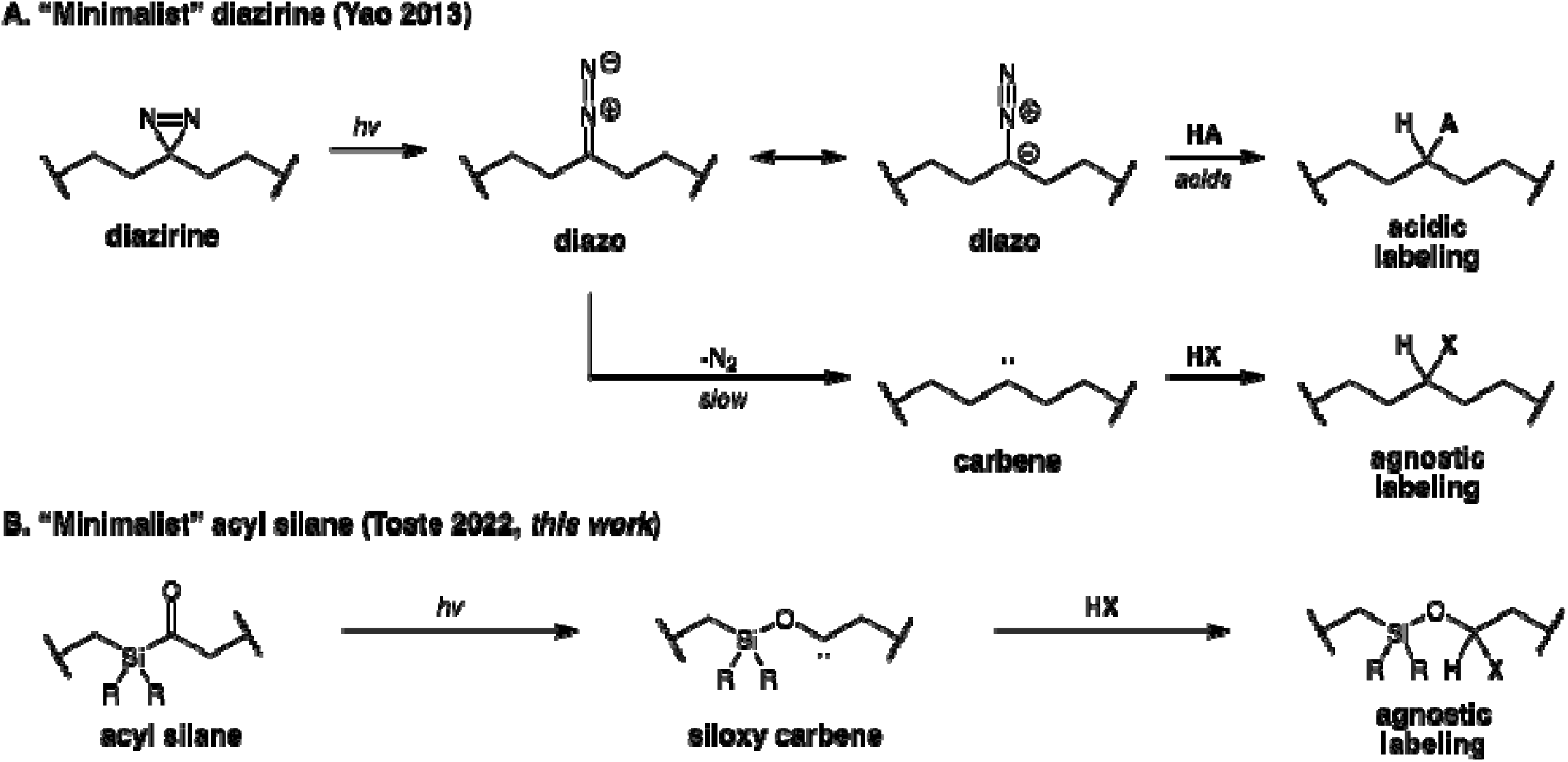
Photoaffinity labeling with diazirines versus acyl silanes. **(A)** Pathways for photoaffinity labeling by aliphatic “minimalist” diazirines. **(B)** Pathway for photoaffinity labeling by acyl silanes.

With these considerations in mind, we recently reported a novel chemical framework for PAL based on the photo-Brook rearrangement of acyl silanes^12^. In this method, photoactivation of the acyl silane species with long-wavelength UV light enables the formation of a reactive α-siloxy carbene species, which can react with amino acid residues in an unbiased manner **(Figure 1B).** We reported a small series of acyl silane probes differentiated by substitution around the photoactivatable moiety, and observed improved labeling profiles as the steric bulk increased. In this initial report, we validated labeling with established small-molecule protein binders in pure proteins and cell lysates. Here, we further optimize our acyl silane reagents and workflow for using these PAL probes for live cell chemoproteomic target identification. We demonstrate that avoiding the diazo pathway leads to a reduction in non-specific protein labeling and then develop a modular synthetic route to enable access to diversified acyl silanes, which can enhance probe properties. Taken together, these findings establish acyl silanes as another alternative to traditional diazirine-based PAL probes for live-cell workflows and highlight the importance of modularity in a PAL probe scaffold.

## Results

### Applying Acyl Silane Photoaffinity Handles in Cellular Chemoproteomic Applications

Our initial report on acyl silane photoaffinity scaffolds aimed to develop a new scaffold for PAL that could avoid the deleterious side reactions associated with diazirines, while also introducing a modular handle through which photo- and physico-chemical properties could be tuned. Through testing several probes, we determined that the labeling efficiency of these probes correlated with the steric bulk about the silane. Additionally, we observed that the bulkier probes were less prone to thermal background reactions, likely resulting from an attack by lysine followed by a thermal Brook rearrangement^12^. Our initial study examined these labeling events in the context of whole proteins and proteomes. We were further interested in evaluating whether these probes could be used in live cells to facilitate target identification in complex biological settings. To assess this possibility, we generated a PAL probe based on the BET inhibitor JQ1, bearing a bulky *i*Pr-substituted acyl silane handle and an alkyne handle, **JQ1-*i*Pr,** for subsequent cellular treatment, photoaffinity-mediated capture of protein targets, copper-catalyzed azide-alkyne cycloaddition (CuAAC) appendage of a biotin-azide enrichment handle, and avidin-enrichment and chemoproteomic identification of targets. We also generated a probe based on the established diazirine PAL handle, **JQ1-DA**, to compare the performance of **JQ1-*i*Pr** with that of an established chemoproteomic probe^1^. Chemoproteomic profiling of both probes in K562 leukemia cells *in situ* showed significant enrichment (p<0.01) of the JQ1 target BRD4 over vehicle-treated control cells. Excitingly, the **JQ1-*i*Pr** probe showed a 3.4-fold enrichment over the control while the **JQ1-DA** probe showed only a 1.6-fold enrichment **(Figure 2A-2B; Table S1; Table S2)**. Moreover, the **JQ1-DA** probe also significantly enriched many off-targets while also not providing substantial enrichment of BRD4. The **JQ1-*i*Pr** probe also significantly enriched a number of non-specific targets—173 other proteins—by more than 2-fold among 1229 proteins quantified. We thus sought to reduce non-specific background labeling further.

**Figure 2.**
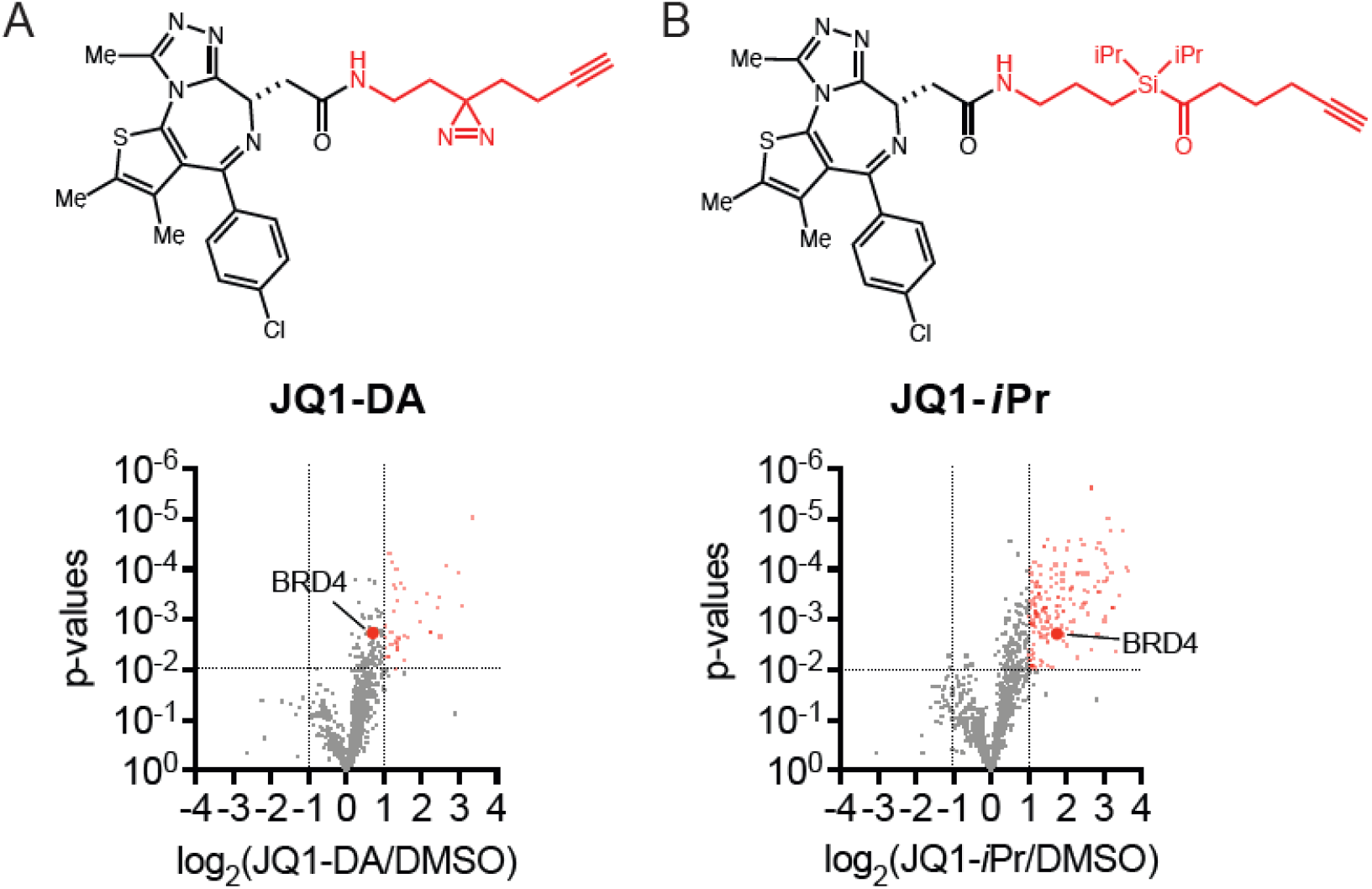
**Chemo**proteomic **profiling with** (+)-JQ1 photoaffinity probes in **live** cells. **(A, B)** K562 cells were treated with DMSO vehicle or PAL probe **JA1-DA (A)** or **JQ1-*i*Pr (B)** (1 μM) for 1 hour and subjected to irradiation at 365 nm for 20 minutes. Subsequent lysates were subjected to CuAAC with an azide-functionalized biotin handle, after which probe-modified proteins were avidin-enriched, eluted, tryptically digested, and analyzed by TMT-based quantitative proteomics. Data shown are ratio of diazirine probe (**DA-JQ1**) **(A)** or diisopropyl acyl silane (**JQ1-*i*Pr**) **(B)** versus DMSO control proteins. Data are from n=3 biologically independent replicates/group. Full dataset can be found in **Tables S1-S2.**

### Synthesis of Second-Generation Acyl Silanes

While previous efforts have investigated varying the linker of the PAL probe to improve its physicochemical properties, we aimed to take advantage of the substitution handle provided by the silane to modulate probe properties, enabling better comparison to current probes while minimizing the distance between the targeting ligand and reactive species to minimize non-specific labeling further. Additionally, a small library of PAL probes that differed only by substitution on the photoactivatable warhead could enable higher-confidence target identification.

As we have established the importance of steric bulk around the acyl silane for labeling efficiency, we hypothesized that a diethyl- or silacyclopentyl-substituted probe could reduce hydrophilicity while preserving labeling efficiency. As such, we selected such dichlorosilanes for our initial reaction screens **(Figure S1).** We began our efforts by examining our previously reported synthetic route to acyl silane probes. In our initial studies, the silane moiety was installed through initial hydrosilylation of an alkene with a disubstituted chlorosilane (HR_2_SiCl). While this route was successful for our initial series of substituents, the relative dearth of these chlorosilane reagents hampered further probe diversification. We therefore turned to the much more available dichlorosilane species for our diversification attempts. The installation of the silane was achieved by reacting the respective dichlorosilane with a Grignard species, followed by the capture of the silyl chloride intermediate by a lithiated dithiane. We found that bromoethyl dioxolane was the most compatible synthon for the Grignard sequence **(Figure S2).** The installation of the alkyne moiety via deprotonation and substitution yielded compound **2**. The tetramethylsilane (TMS) functionality on the alkyne was deprotected by addition of potassium carbonate in methanol to give terminal alkynes **3**. Sequential acid-promoted deprotection of the acetal and reduction of the corresponding aldehyde with sodium borohydride afforded alcohol intermediates **4**. Following installation of the phenyl carbamate moiety, deprotection of dithianes **5** by PIFA revealed substituted probes **15-Et** and **15-cyclo (Figure 3).**

**Figure 3.**
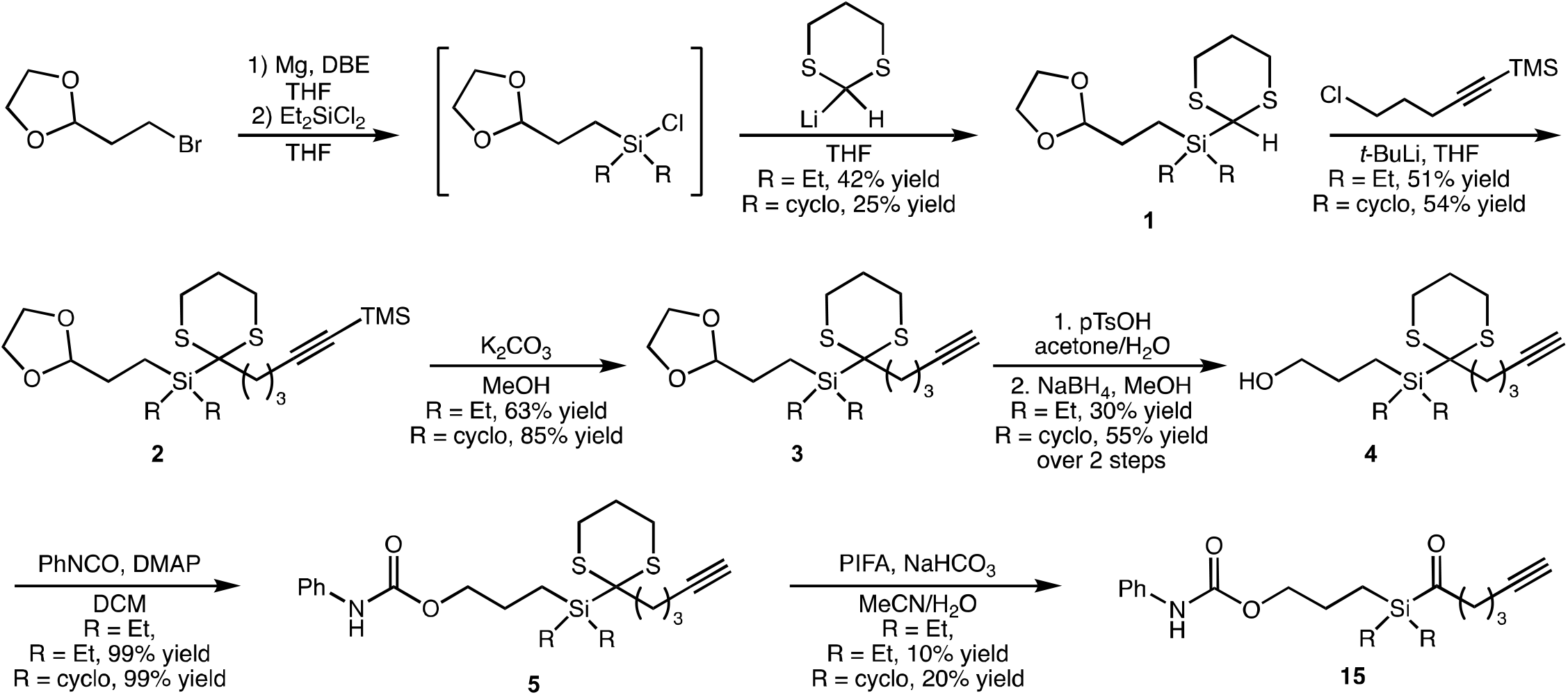
Synthesis of proof-of-concept photoaffinity probes containing diversified silane substitution.

### Evaluation of Second-Generation Acyl Silane Probes

With two new substituted acyl silane probes in hand, we sought to evaluate their labeling compared to the best performing acyl silane from our original report, the isopropyl-substituted probe **15-*i*Pr**. We began by evaluating their labeling performance in cell lysate. Upon photoaffinity labeling in cell lysates and CuAAC-mediated appendage of an azide-functionalized rhodamine handle, labeled proteins were subsequently visualized by in-gel fluorescence. Probe **15-Et** displayed light-dependent labeling of proteins to a similar extent as probe **15-*i*Pr** while preserving the minimal thermal background labeling as its bulkier congener **(Figure 4).** However, samples treated with silacyclopentane-substituted probe **15-cyclo** showed significantly greater fluorescence in the samples incubated in the absence of light, suggesting the susceptibility of this species towards thermal background reactivity **(Figure 4).** From these studies, we determined that the ethyl substituted probe was better suited than the cyclopentane substituted probe for further design of chemoproteomic probes.

**Figure 4.**
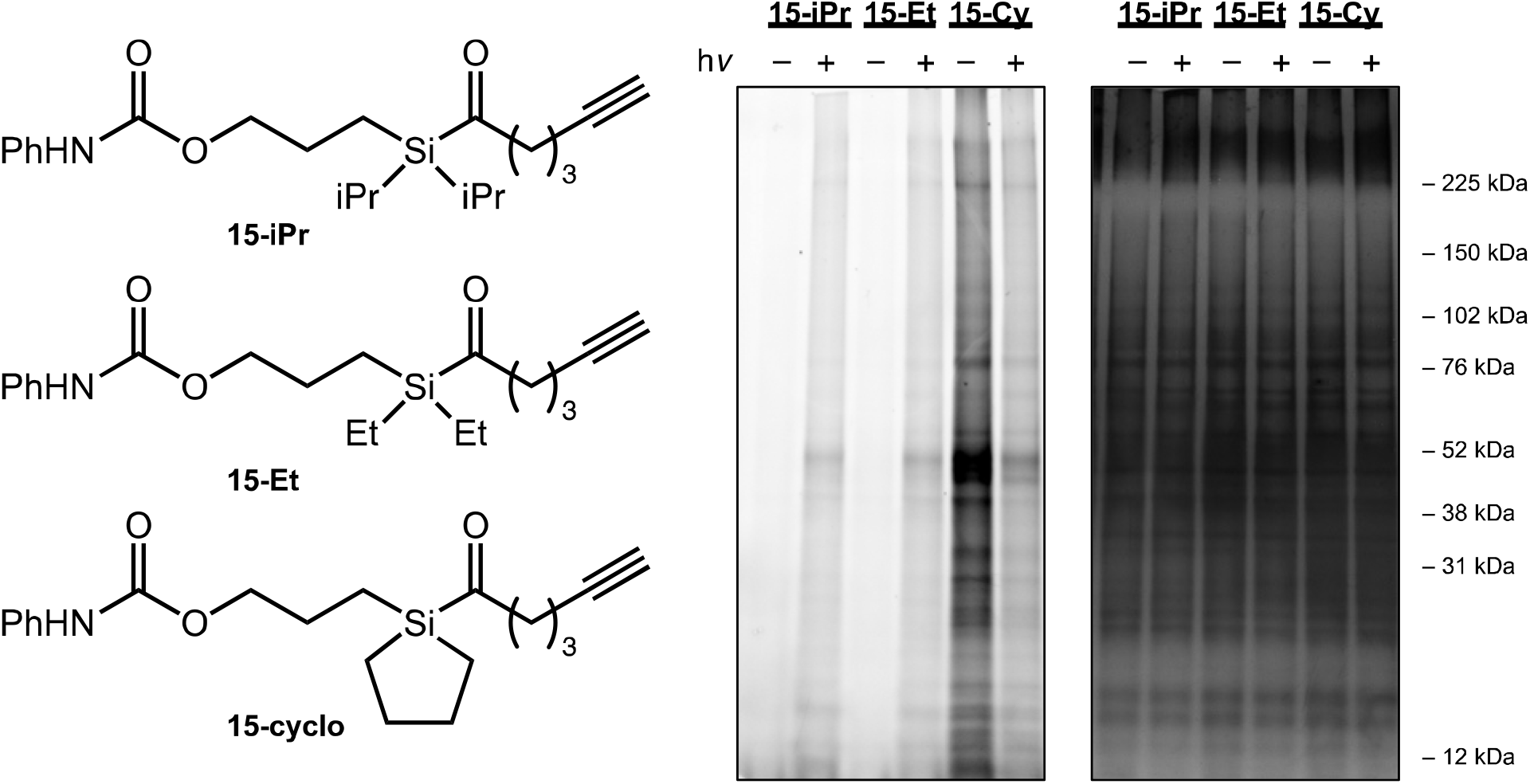
Evaluation of labeling by photoaffinity probes. Evaluation of labeling by proof-of-concept photoaffinity probes (15-iPr, 15-Et, and 15-cyclo) in K562 cell lysate by in-gel fluorescence (left image) following Cu click reaction with Rh–N_3_ after irradiation at 365 nm or incubation in the dark. Protein loading was confirmed following silver staining (right image). Gels are representative from n=3 biologically independent replicates per group.

**Figure 5.**
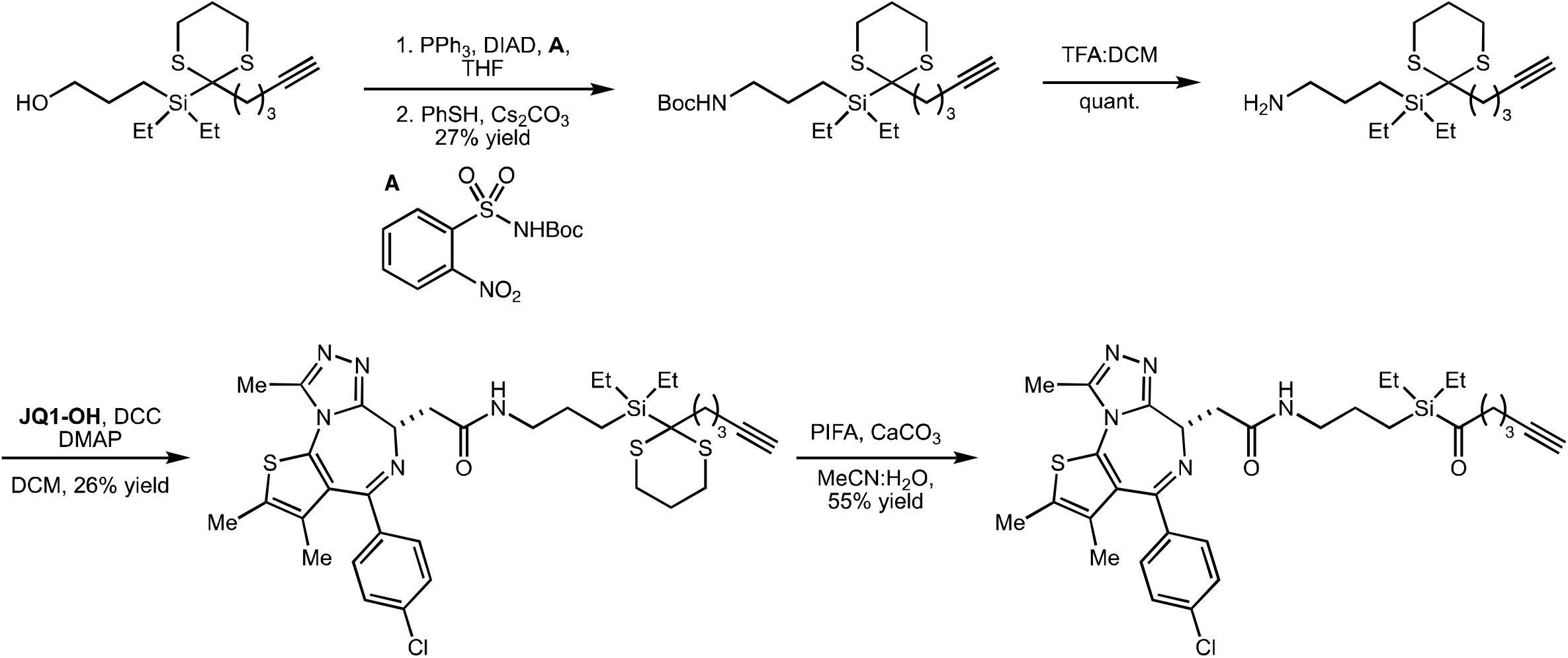
Synthetic route towards fully-functionalized probe JQ1-Et.

### Synthesis and Testing of Fully-Functionalized Second-Generation Acyl Silane Probes

To validate our hypothesis that reduced hydrophobicity could lead to improved labeling specificity, we next synthesized the (+)-JQ1-ligated probe **JQ1-Et**. Conversion of the alcohol **4-Et** to amine **6-Et** under Mitsunobu conditions enabled access to versatile amine handle **7-Et** after removal of the Boc protecting group. An amide coupling of the carboxylic acid derived from (+)-JQ1 to this amine allowed access to the fully-functionalized diethyl acyl silane **JQ1-Et (Figure 6A; Table S3).** Cellular chemoproteomic profiling of **JQ-Et** not only demonstrated significant enrichment of BRD4, but also less non-specific labeling and enrichment with 58 other proteins enriched out of 683 total quantified proteins compared to 173 proteins with **JQ1-*i*Pr (Figure 6A; Table S4).** Comparing the proteins enriched significantly by **JQ1-iPr** and **JQ1-Et**, 25 proteins were enriched significantly by both probes, one of which was BRD4 **(Figure 6B, Table S4).** Overall, our findings support our hypothesis that reduced hydrophobicity of the second-generation probe reduces non-specific labeling and enrichment of proteins.

**Figure 6.**
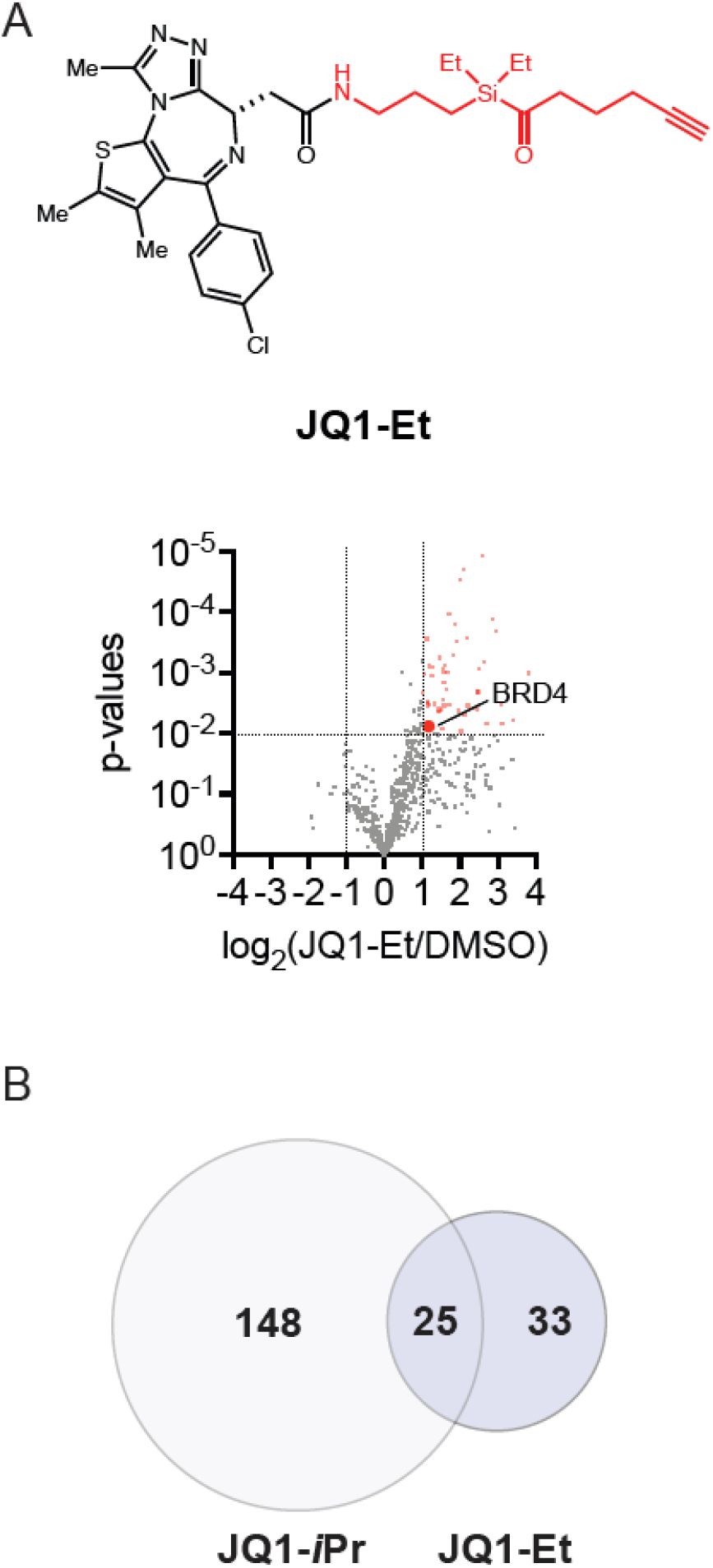
Chemoproteomic profiling of JQ1-Et. **(A)** Quantitative proteomics of (+)-JQ1 diethyl acyl silane probe (**JQ1-Et**) in whole cells. K562 cells were treated with DMSO vehicle or probe (1 μM) for 1 hour and subjected to irradiation at 365 nm for 20 minutes. Subsequent lysates were subjected to CuAAC with an azide-functionalized biotin handle, after which probe-modified proteins were avidin-enriched, eluted, and digested, and analyzed by TMT-based quantitative proteomics. Data shown are ratio of **JQ1-Et** probe vs. DMSO control enriched proteins. Data are from an n=3 biologically independent replicates per group. Full dataset can be found in **Table S3. (B)** Venn diagram showing overlap in groups of proteins significantly enriched (p-value < 0.01) by either acyl silane probe **JQ1-*i*Pr** or **JQ1-Et**, which only differ by substitution about the silane. BRD4 was found in this dataset; full list of overlapping proteins is included in **Table S4**.

In testing our new generation of acyl silane probes in live cells, we aimed to facilitate peptide mapping to observe the direct modification of BRD4 with our probes and explore potential residue selectivity, as has been achieved by previous groups with diazirine handles^5^. However, we were unable to identify any mass adducts corresponding to our probes in proteomics experiments. We surmised that the siloxy product of the acyl silane PAL reaction could be acid-labile, as our LC-MS/MS workflows require 0.1% formic acid in the mobile phase. To evaluate this, we performed PAL labeling in lysate using a modified workflow that included titration of formic acid after labeling and the click reaction, but before adding the reducing reagent and running SDS/PAGE. As expected, the amount of labeling product observed decreased with increased acid **(Figure S3).** The acid-lability of these probes is under further investigation in our labs for use as a traceless probe for protein labeling. In total, our work demonstrates the utility of acyl silane PAL probes in cellular chemoproteomic workflows.

## Discussion

Photoaffinity labeling coupled with chemoproteomic profiling is a powerful tool for the deconvolution of small molecule-protein interactions, but its use in live cell applications is hampered by the non-specific labeling observed with commonly used alkyl diazirine crosslinkers. We recently reported acyl silanes as a novel scaffold for photoaffinity labeling, based on the 1,2-photo-Brook rearrangement of acyl silanes to form an α-siloxy carbene intermediate. In this study, we optimize and demonstrate the utility of acylsilanes for PAL and cellular chemoproteomic and target identification applications as exemplified with BRD4 probes. To improve probe labeling properties, we developed a modular synthetic route to enable access to acyl silanes with diverse substituents at silicon and demonstrated that reducing the hydrophobicity of the photoaffinity probe improves specificity of probe labeling in living cells. We do, however, still observe some non-specific labeling and enrichment of proteins that are unlikely targets of JQ1. Specificity of labeling in our study could perhaps be further increased by comparison of additional linker types functionalized with either acyl silane. We also showed that site of labeling was not possible with our current scaffolds due to their acid liability under mass spectrometry conditions. Future work may include optimization of adduct stability under acidic conditions. Alternatively, these acid-labile adducts could allow for temporal control of covalent labeling, which can be released in a scarless manner via acidification of the samples. Overall, our results establish acyl silanes as a new tool for photoaffinity labeling for target identification studies.

## Acknowledgment

We thank the members of the Nomura Research Group and Novartis BioMedical Research for their critical review of the manuscript. This work was also supported by Novartis Biomedical Research, the National Science Foundation Molecular Foundations for Biotechnology (MFB) grant (2127788), and the National Institutes of Health (R35CA263814, R01CA240981, UM1CA29410). We also thank Dr. Hasan Celik, Lund, and the UC Berkeley NMR facility in the College of Chemistry (CoC-NMR) for spectroscopic assistance. Instruments in the College of Chemistry NMR facility are partly supported by NIH S10OD024998. We also thank the Quantitative Biosciences Institute (Dr. Robert Maxwell). ACSP would like to thank Deborah Zhuang for assistance with high-resolution mass spectrometry.

## Competing Financial Interests Statement

DKN is a co-founder, shareholder, and scientific advisory board member for Frontier Medicines and Zenith. DKN is also on the scientific advisory board of The Mark Foundation for Cancer Research, Photys Therapeutics, Oerth Bio, Apertor Pharmaceuticals, and Ten30 Biosciences. DKN is also an Investment Advisory Partner for a16z Bio, an Advisory Board member for Droia Ventures, and an iPartner for The Column Group.

