## Supporting Information for "Development of Second-Generation Acyl Silane Photoaffinity Probes for Cellular Chemoproteomic Profiling"

**List of supplementary tables**

**Table S1.**
TMT proteomics data quantifying proteins enriched by probe **JQ1-DA** vs DMSO control.

**Table S2.**
TMT proteomics data quantifying proteins enriched by probe **JQ1-iPr** vs DMSO control.

**Table S3.**
TMT proteomics data quantifying proteins enriched by probe **JQ1-Et** vs DMSO control.

**Table S4.**
Merged TMT proteomics data showing only proteins enriched by both probes **JQ1-iPr** and **JQ1-Et** compared to DMSO by at least 2-fold and p-value < 0.05 across n=3 biologically independent replicates/group.

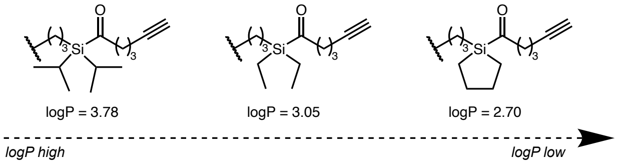


**Figure S1.** Calculated logP of acyl silane fragments.

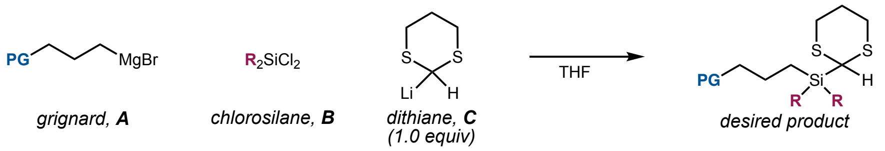


| PG (equiv A) | R (equiv B) | Order of addition | Yield |
|---|---|---|---|
| OTBS (1.2) | Et (1.1) | **B** to **C**, add to **A** | 0% |
| OTHP (2.0) | Et (1.0) | **C** to **B**, add to **A** | 13% |
| OTHP (1.5) | Et (1.2) | **A** to **B**, add to **C** | 0% |
| OBn (1.5) | Et (1.2) | **A** to **B**, add to **C** | 52% |
| dioxolane (1.5) | Et (1.2) | **A** to **B**, add **C** @ -78 | 42% |
| dioxolane (1.5) | silacyclopentane (1.2) | **A** to **B**, add **C** @ -78 | 25% |

**Figure S2.** Condition screen for Grignard/lithiation sequence.

Commented [AP1]: Delete?

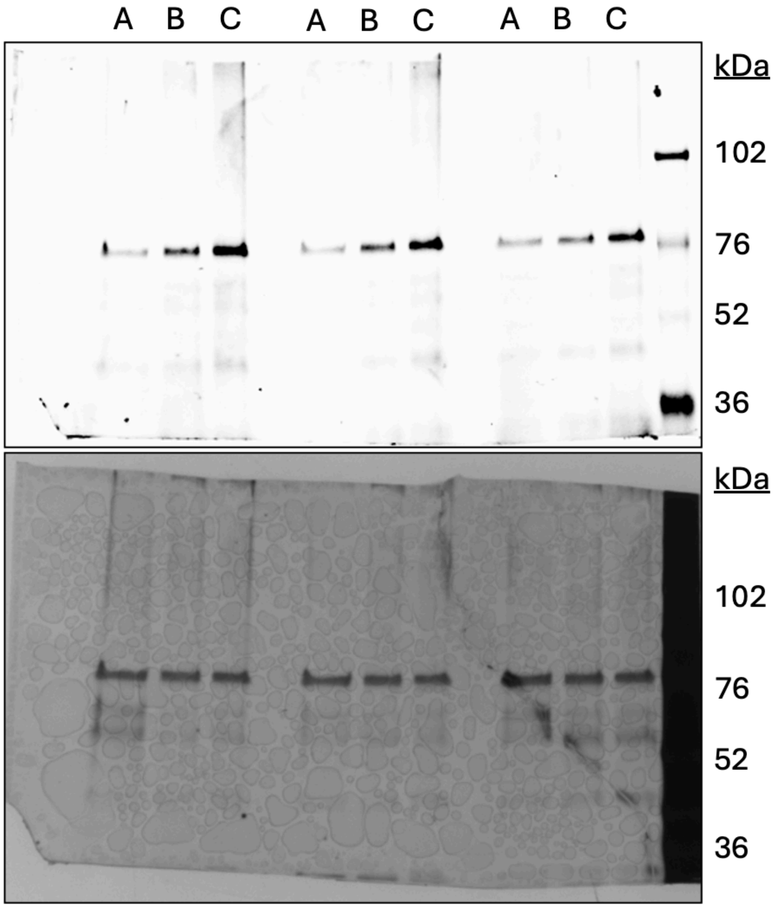


**A: 0.1% FA, B: 0.01% FA, C: 0% FA.**

**Figure S3. Acidic conditions attenuate JQ1-Et probe labeling of recombinant BRD4.** Recombinant BRD4 containing only BD1 and BD2 was treated with **JQ1-Et**, irradiated, and functionalized with rhodamine azide using CUAAC. Samples were treated for 1 hour with either 0.1%, 0.01% or 0% formic acid, then neutralized with sodium hydroxide solution and analyzed via SDS-PAGE. Top image: rhodamine labeling; bottom image: silver staining.

**Materials and Methods**

**Cell culture**

K562 cells were obtained from the UC Berkeley Cell Culture Facility and were cultured in RPMI medium containing 10% (v/v) fetal bovine serum (FBS) and maintained at 37°C with 5% $CO_2$.

**Preparation of cell lysate**

Cells were pelleted by centrifugation (1400 xg, 5 minutes, 4 °C). Pellets were resuspended in RIPA buffer containing protease inhibitor and lysed on ice, and clarified by centrifugation (12,000 xg, 10 minutes, 4 °C). Lysate was transferred to low-adhesion microcentrifuge tubes. Proteome concentrations were determined using BCA assay and lysate was diluted with PBS to appropriate working concentrations.

**Gel-based ABPP in cell lysate**

K562 cell lysate (1.0 µg/µL) in pH 7.4 PBS (50 µL) was treated with a prepared stock solution of DMSO or PAL probe (10 µM) in DMSO and incubated in the dark for 30 minutes at RT. Samples were irradiated for 30 minutes on ice at 365 nm. CuAAC was performed to append rhodamine-azide (1 µM final concentration) onto alkyne probe-labeled proteins. Samples were then diluted with 4x reducing Laemmli SDS sample loading buffer and heated at 95 °C for 5 minutes. The samples were separated on precast 4-20% Criterion TGX gels (Bio-Rad). Prior to analysis, gels were fixed in a solution of 10% acetic acid and 30% ethanol for 1 hour. Probe-labeled proteins were analyzed by in-gel fluorescence using a ChemiDoc MP (Bio-Rad). Protein loading was assessed by silver stain.

**Gel-based ABPP in whole cells**

K562 cells were treated with either DMSO vehicle or PAL probe (10 µM) for 1 hour in serum-free media and subsequently irradiated for 20 minutes at 365 nm. Lysate was prepared as described above. CuAAC was performed to append rhodamine-azide (1 µM final concentration) onto alkyne probe-labeled proteins. Samples were then diluted with 4x reducing Laemmli SDS sample loading buffer and heated at 95 °C for 5 minutes. The samples were separated on precast 4-20% Criterion TGX gels (Bio-Rad). Prior to analysis, gels were fixed in a solution of 10% acetic acid and 30% ethanol for 1 hour. Probe-labeled proteins were analyzed by in-gel fluorescence using a ChemiDoc MP (Bio-Rad). Protein loading was assessed by silver stain.

**Acid Cleavage Gel-based ABPP with Recombinant BRD4**

Recombinant BRD4 (BD1+BD2, purchased from Cayman Chemical, C833Y26) was diluted to 0.02 µg/µL (0.3 µM) in 1X PBS and treated with 10 µM JQ1-Et for 30 minutes at room temperature and subsequently irradiated for 20 minutes at 365 nm. CuAAC was performed to append rhodamine-azide (1 µM final concentration) onto alkyne probe-labeled proteins. Samples were adjusted to 0.1%, 0.01%, or 0% formic acid and incubated for 1 hour, then neutralized with an equal molar ratio of sodium hydroxide solution. Samples were then diluted with 4x reducing Laemmli SDS sample loading buffer and heated at 95 °C for 5 minutes. The samples were separated on precast 4-20% Criterion TGX gels (Bio-Rad). Prior to analysis, gels were fixed in a solution of 10% acetic acid and 30% ethanol for 1 hour. Probe-labeled BRD4 was analyzed by in-gel fluorescence using a ChemiDoc MP (Bio-Rad). Protein loading was assessed by silver stain.

**TMT proteomics**

K562 cells at 80% confluency were treated with DMSO vehicle, **JQ1-iPr** (1 µM), or **JQ1-DA** (1 µM). After 1 hour of incubation, the cells were irradiated on ice for 30 minutes, harvested, and lysed via sonication. Each lysate was normalized to a concentration of 2 mg/mL, and 500 µL of each lysate was transferred to a separate tube. To each tube containing 500 µL of cell lysate, 10 µL of 20 mM biotin picolyl azide (Sigma-Aldrich, 900912) in DMSO, 10 µL of 50 mM TCEP in $H_2O$,10 µL of 50 mM $CuSO_4$ in $H_2O$, and 30 µL of TBTA ligand (1.3 mg/mL in 1:4 DMSO/*t*-BuOH, Cayman Chemical, 18816) were added. The reaction mixture was incubated at room temperature for 60 minutes with agitation. The reaction was quenched by protein precipitation. After washing the protein pellets twice with cold MeOH (4 °C), the pellets were redissolved in 250 µL of 1.2% SDS/PBS (w/v). The samples were heated at 95 °C for 5 minutes and then spun at 6500 xg for 5 minutes. The supernatant was transferred to new tubes containing 1.25 mL of PBS was added to the remaining sample to reduce the total SDS concentration to less than 0.2% SDS/PBS (w/v). 170 µL of streptavidin agarose beads (ThermoFisher, 20353) were added to the lysate-containing tubes, and the samples were incubated at 4 °C on a rotator overnight. After incubation, the samples were brought to 23 °C, and the beads were spun down in a centrifuge (1300 xg, 2 minutes). The supernatant was removed, and the beads were washed three more times with 500 µL of PBS and

500 µL of $H_2O$. The beads were resuspended in 50 µL of 0.1% SDS/PBS (w/v) and the samples were further treated with DTT (90 µg per 1 mg protein) for 20 minutes at 65 °C. Iodoacetamide (237 µg per 1 mg protein) was added and the samples were incubated for 1 hour at 37 °C. After incubation, 25.8 µL of a premixed solution containing 25 µL 100 mM triethylammonium biocarbonate (TEAB), 0.25 µL 100 mM $CaCl_2$, and 0.5 µL 20 mg/mL trypsin (Promega V5111) was added to each sample. The samples were incubated horizontally overnight at 37 °C with agitation. The collected peptide-containing samples were dried using a vacufuge and resuspended in 100 µL of 50 mM TEAB before labeling using commercially available TMT10plex tags (ThermoFisher, PN 90110). After labeling, 30 µg of each labeled sample was combined and dried using a vacufuge. Dried samples were redissolved with 300 µL of 0.1% TFA in $H_2O$ and further fractionated using high-pH reversed-phase peptide fractionation kits (ThermoFisher, PN 84868) following the manufacturer's protocol.

Mass spectrometry analysis was performed on an Orbitrap Eclipse Tribrid Mass Spectrometer with a High Field Asymmetric Waveform Ion Mobility (FAIMS Pro) Interface (Thermo Scientific) with an UltiMate 3000 Nano Flow Rapid Separation LCnano System (Thermo Scientific). Off-line fractionated samples (5 µl aliquot of 15 µl sample) were injected via an autosampler (Thermo Scientific) onto a 5 µl sample loop which was subsequently eluted onto an Acclaim PepMap 100 C18 HPLC column (75 µm x 50 cm, nanoViper). Peptides were separated at a flow rate of 0.3 µl/min using the following gradient: 2 % buffer B (100% acetonitrile with 0.1% formic acid) in buffer A (95:5 water:acetonitrile, 0.1% formic acid) for 5 minutes, followed by a gradient from 2 – 40 % buffer B from 5 – 159 minutes, 40 – 95% buffer B from 159 to 160 minutes, holding at 95% B from 160 – 179 minutes, 95 – 2% buffer B from 179 – 180 minutes, and then 2% buffer B from 180 – 200 minutes. Voltage applied to the nano-LC electrospray ionization source was 2.1 kV. Data was acquired through an MS1 master scan (Orbitrap analysis, resolution 120,000, 400-1800 m/z, RF lens 30%, heated capillary temperature 250 °C) with dynamic exclusion enabled (repeat count 1, duration 60 s). Data-dependent data acquisition comprised a full MS1 scan followed by sequential MS2 scans based on 2 s cycle times. FAIMS compensation voltages (CV) of -35, -45, and -55 were applied. MS2 analysis consisted of: quadrupole isolation window of 0.7 m/z of precursor ion followed by higher energy collision dissociation (HCD) energy of 38% with an orbitrap resolution of 50,000.

Acquired MS data was processed using ProLuCID search methodology in IP2 v.3-v.5 (Integrated Proteomics Applications, Inc.). Trypsin cleavage specificity (cleavage at K, R except if followed by P) allowed for up to 2 missed cleavages. Carbamidomethylation of cysteine was set as a fixed modification, methionine oxidation, and TMT-modification of N-termini and lysine residues were set as variable modifications. Reporter ion ratio calculations were performed using summed abundances with the most confident centroid selected from the 10 ppm window. Only peptide-to-spectrum matches that are unique assignments to a given identified protein within the total dataset and with an average intensity of over 40,000 are considered for protein quantitation. High confidence protein identifications were reported with a <1% false discovery rate (FDR) cut-off. Differential abundance significance was estimated using ANOVA with Benjamini-Hochberg correction to determine p-values.

**GO analysis**
Gene ontology (GO) analysis for subcellular localization of labeled proteins was performed using the DAVID Gene Functional Classification Tool at https://david.ncifcrf.gov/summary.jsp.

**LogP calculation**
Calculation of predicted logP values were performed using the Molinspiration Cheminformatics free web services at https://www.molinspiration.com.

**General Synthetic Methods**
Unless otherwise noted, all reagents were purchased from commercial suppliers and used without further purification. Tetrahydrofuran, dichloromethane, diethyl ether, toluene, trimethylamine, and *N,N*-dimethylformamide were purified by passage through an activated alumina column under argon. Thin-layer chromatography (TLC) analysis of reaction mixtures was performed using Merck silica gel 60 F254 TLC plates and visualized under UV or by staining with $KMnO_4$ or *p*-anisaldehyde. Column chromatography was performed on Merck Silica Gel 60 Å, 230 X 400 mesh. Nuclear magnetic resonance (NMR) spectra were recorded using Bruker AV-600, AV-500, Neo-500, Neo-501, AVB-400, and AVQ-400 spectrometers at the Pines Magnetic Resonance Center's Core NMR Facility at the University of California, Berkeley. All chemical shifts are reported in the standard notation of δ parts per million relative to the residual solvent peak at 7.26 ($CDCl_3$) or 3.31 ($CD_3OD$) for $^1$H and 77.16 ($CDCl_3$) or 49.00 ($CD_3OD$) for $^{13}$C as an internal reference. Splitting patterns are indicated as follows: s = singlet, d = doublet, t = triplet, q = quartet, p = pentet, hept = heptet/septet, m = multiplet, br = broad, dt = doublet of triplet resonance. These NMR instruments at UC Berkeley are funded in part by the NIH (S10OD024998). High-resolution mass spectral analyses (ESI-MS) were carried out using an Agilent 1260 series LC equipped with a 6530 series Q-TOF (ESI) located in the Toste group at the University of California, Berkeley or obtained from the QB3 mass spectral facility at the University of California, Berkeley on a Thermo LTQ-FTICR (7T, ESI). Solvent abbreviations are reported as follows: EtOAc = ethyl acetate, hex = hexanes, DCM = dichloromethane, $Et_2O$ = diethyl ether, MeOH = methanol, THF = tetrahydrofuran, DMSO = dimethylsulfoxide, $Et_3N$ = trimethylamine, MeCN = acetonitrile.

**Synthetic Methods and Characterization**
**Synthesis of acyl silane 15-Et**

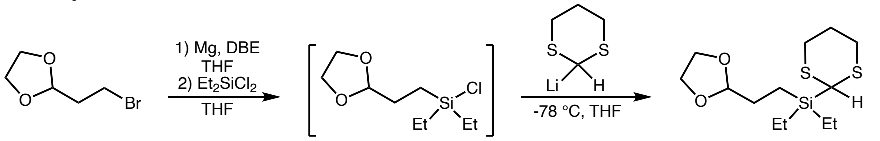


**(2-(1,3-dioxolan-2-yl)ethyl)(1,3-dithian-2-yl)diethylsilane (1-Et)**
A flame-dried three-neck flask fitted with a reflux condenser and cooled under $N_2$ was charged with magnesium turnings (0.6321 g, 26 mmol, 2.03 equiv.) and THF (20 mL). A small amount of 1,2-dibromoethane was added and the solution was heated with a heat gun until ethylene bubbles were observed. The flask was then placed in a water bath and dioxolane (2.35 mL, 20 mmol, 1.56 equiv.) was added slowly. The mixture was allowed to stir at 40 °C until a majority of the magnesium was consumed (ca. 2 hours). The resulting Grignard reagent was added via cannula to a separate flame-dried flask containing dichlorodiethylsilane (2.25 mL, 15.4 mmol, 1.20 equiv.) and THF (10 mL) at 0 °C. The mixture was allowed to warm to room temperature over 2 hours. Meanwhile, a separate round bottom flask cooled under vacuum was charged with 1,3-dithiane (1.54 g, 12.82 mmol, 1.0 equiv.) and THF (25 mL). The reaction vessel was cooled to –78 ˚C and a solution of *n*BuLi (2.5 M in hexanes, 5.64 mL) was added dropwise resulting in a persistent, but faint yellow color. The flask was allowed to warm to room temperature over 30 minutes and then cooled back to –78 °C and transferred via cannula to the flask containing silyl chloride at –78 °C. The solution was allowed to warm to room temperature overnight. After 16 hours, the reaction was diluted with EtOAc, quenched by addition of sat. aqueous $NH_4Cl$ and washed with water. The combined aqueous layers were then extracted by EtOAc 2X and the combined organic layers were washed with brine, dried over $MgSO_4$, filtered, and concentrated under reduced pressure. The crude oil was then purified via flash-column chromatography (gradient elution 0% – 10% EtOAc/Hex) to give **1-Et** as a clear oil (1.646 g, 5.369 mmol, 42% yield).
**$^1$H NMR** (600 MHz, $CDCl_3$): δ 4.83 (t, *J* = 4.6 Hz, 1H), 4.00 – 3.93 (m, 2H), 3.90 – 3.83 (m, 2H), 3.82 (s, 1H), 2.87 (ddd, *J* = 14.6, 12.6, 2.6 Hz, 2H), 2.70 (ddd, *J* = 14.0, 4.2, 2.9 Hz, 2H), 2.11 (dtt, *J* = 14.1, 4.7, 2.7 Hz, 1H), 2.07 – 1.97 (m, 1H), 1.75 (dt, *J* = 13.4, 4.5 Hz, 2H), 1.02 (t, *J* = 7.9 Hz, 6H), 0.83 – 0.76 (m, 2H), 0.72 (q, *J* = 7.9 Hz, 4H). **$^{13}$C NMR** (151 MHz, $CDCl_3$): δ 106.12, 65.10, 32.30, 31.55, 28.20, 26.51, 7.53, 4.29, 2.60. **HRMS** (ESI): calculated for $[C_{13}H_{26}O_2S_2Si+H]^+$ required *m/z* 307.1216, found *m/z* 307.1223.

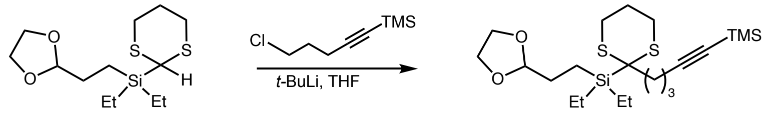

**(2-(1,3-dioxolan-2-yl)ethyl)diethyl(2-(5-(trimethylsilyl)pent-4-yn-1-yl)-1,3-dithian-2-yl)silane (2-Et)**

A flame-dried round bottom flask cooled under vacuum was charged with **5.1-Et** (1.218 g, 3.97 mmol, 1.0 equiv.) and THF (20 mL, 0.2 M) and cooled to –78 °C, followed by the dropwise addition of *t*BuLi (1.6 M in pentane, 2.61 mL, 1.05 equiv.). After stirring for 30 minutes at –78 °C, the solution was allowed to warm to room temperature over 30 minutes and then cooled back down to –78 °C. Next, (5-chloropent-1-yn-1-yl)trimethylsilane (0.78 mL, 4.37 mmol, 1.1 equiv.) was added dropwise and the reaction was allowed to warm to room temperature overnight. After 16 hours, the reaction was quenched by the addition of sat. aqueous $NH_4Cl$ and extracted with EtOAc. The combined organic layers were washed with saturated brine, dried over $MgSO_4$, and concentrated to an oil. Purification of the crude material by flash column chromatography (5% EtOAc/hex) yielded the desired product **2-Et** as a clear, colorless oil (0.896 g, 2.014 mmol, 51% yield).

**$^1$H NMR** (600 MHz, $CDCl_3$): δ 4.84 (t, *J* = 4.6 Hz, 1H), 4.00 – 3.93 (m, 2H), 3.89 – 3.83 (m, 2H), 3.10 (ddd, *J* = 14.8, 12.7, 2.7 Hz, 2H), 2.39 (dt, *J* = 13.7, 3.8 Hz, 4H), 2.32 (t, *J* = 6.5 Hz, 2H), 2.07 – 1.99 (m, 1H), 1.93 (qt, *J* = 12.9, 3.3 Hz, 1H), 1.83 (ddd, *J* = 9.4, 8.2, 4.5 Hz, 2H), 1.76 – 1.67 (m, 2H), 1.06 (t, *J* = 7.9 Hz, 6H), 0.92 – 0.87 (m, 2H), 0.84 – 0.77 (m, 4H), 0.15 (s, 9H). **$^{13}$C NMR** (151 MHz, $CDCl_3$): δ 107.32, 106.34, 85.23, 65.08, 39.57, 36.41, 28.88, 26.86, 25.32, 23.49, 20.26, 8.27, 4.95, 3.28, 0.30, -1.04. **HRMS** (ESI) calculated for $[C_{21}H_{40}O_2S_2Si_2+H]^+$ required *m/z* 445.2081, found *m/z* 445.2112.

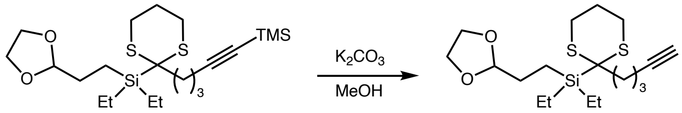


**(2-(1,3-dioxolan-2-yl)ethyl)diethyl(2-(pent-4-yn-1-yl)-1,3-dithian-2-yl)silane (3-Et)**

A scintillation vial was charged with **2-Et** (2.160 g, 4.856 mmol, 1.0 equiv.), potassium carbonate (2.013 g, 14.567 mmol, 3 equiv.), and MeOH (12 mL). The reaction was allowed to stir at room temperature for 2 hours and was then diluted with ether and washed with sat. aqueous $NH_4Cl$, water, and brine. The organic layers were dried over $MgSO_4$, filtered, and concentrated under reduced pressure. The crude oil was purified using flash column chromatography (10% EtOAc/hex) to give alkyne **3-Et** (1.139 g, 3.056 mmol, 63% yield) as a clear oil.

**$^1$H NMR** (500 MHz, $CDCl_3$) δ 4.84 (t, *J* = 4.7 Hz, 1H), 4.01 – 3.92 (m, 2H), 3.91 – 3.80 (m, 2H), 3.07 (ddd, *J* = 15.0, 12.7, 2.7 Hz, 2H), 2.39 (ddd, *J* = 13.3, 6.8, 3.1 Hz, 4H), 2.27 (td, *J* = 6.8, 2.7 Hz, 2H), 2.08 – 2.00 (m, 1H), 1.99 (t, *J* = 2.6 Hz, 1H), 1.92 (qt, *J* = 12.9, 3.2 Hz, 1H), 1.83 (ddd, *J* = 9.4, 8.2, 4.5 Hz, 2H), 1.79 – 1.67 (m, 2H), 1.05 (t, *J* = 7.9 Hz, 6H), 0.92 – 0.85 (m, 2H), 0.81 (q, *J* = 8.0 Hz, 4H). **$^{13}$C NMR** (126 MHz, $CDCl_3$): δ 106.32, 84.38, 68.94, 65.08, 39.42, 36.71, 28.87, 26.86, 25.27, 23.57, 18.92, 8.26, 4.90, 3.30. **HRMS** (ESI) calculated for $[C_{18}H_{32}O_2S_2Si+H]^+$ required *m/z* 373.1686, found *m/z* 373.1699.

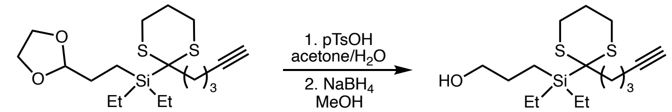


**3-(diethyl(2-(pent-4-yn-1-yl)-1,3-dithian-2-yl)silyl)propan-1-ol (4-Et)**

A 50 mL round-bottom flask was charged with alkyne **3-Et** (1.139 g, 3.056 mmol, 1.0 equiv.) and acetone (40 mL). Water (10 mL) and *p*-TsOH (3.157 g, 18.336 mmol, 6.0 equiv.) were added and the reaction was stirred at room temperature. After 16 hours, the reaction was quenched by addition of sat. aqueous $NaHCO_3$ and extracted with EtOAc. The combined organic layers were washed with brine, dried over $MgSO_4$, filtered, and concentrated to give the aldehyde as a yellow oil which was used immediately in the next step without further purification. A scintillation vial was then charged with aldehyde and MeOH (12.25 mL). The vial was cooled to 0 °C and sodium borohydride (97 mg) was added and the reaction was allowed to stir. After one hour, an additional dose of sodium borohydride (60 mg) was added and the reaction was stirred for an additional hour. The reaction was then quenched by addition of water and extracted with EtOAc. The combined organic layers were washed with brine, dried over $MgSO_4$, filtered, and concentrated under reduced pressure. The crude oil was purified using flash column chromatography (25% EtOAc/hex) to give alcohol **4-Et** (0.294 g, 0.889 mmol, 30% yield over two steps) as a clear oil.

**$^{1}$H NMR** (500 MHz, $CDCl_3$) δ 3.63 (t, *J* = 6.6 Hz, 2H), 3.08 (ddd, *J* = 15.2, 12.7, 2.7 Hz, 2H), 2.40 (ddd, *J* = 13.0, 6.7, 3.0 Hz, 4H), 2.28 (td, *J* = 6.8, 2.6 Hz, 2H), 2.08 – 2.02 (m, 1H), 2.00 (t, *J* = 2.6 Hz, 1H), 1.97 – 1.85 (m, 1H), 1.79 – 1.69 (m, 4H), 1.06 (t, *J* = 7.9 Hz, 6H), 0.85 – 0.78 (m, 6H). **$^{13}$C NMR** (126 MHz, $CDCl_3$) δ 68.98, 66.02, 36.66, 27.72, 26.89, 25.29, 23.55, 18.92, 14.28, 8.31, 6.85, 3.37 (*one resonance not observed)*. **HRMS** (ESI) calculated for $[C_{16}H_{30}OS_2Si+H]^+$ required *m/z* 331.1580, found *m/z* 331.1613.

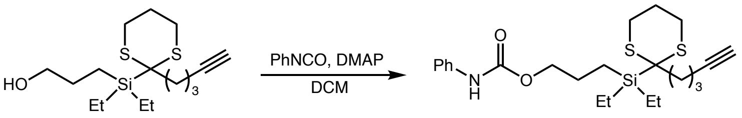


**3-(diethyl(2-(pent-4-yn-1-yl)-1,3-dithian-2-yl)silyl)propyl phenylcarbamate (5-Et)**
A scintillation vial was charged with alcohol **4-Et** (0.050 g, 0.151 mmol, 1.0 equiv.), DMAP (4 mg, 0.032 mmol, 0.2 equiv.), and DCM (0.80 mL). Phenylisocyanate (20 uL, 0.17 mmol, 1.1 equiv.) was added and the reaction was allowed to stir at room temperature overnight. After 16 hours, the reaction was diluted with DCM, washed successively with sat. aqueous $NH_4Cl$, water, and brine, and the organic layer was dried over $MgSO_4$, filtered, and concentrated under reduced pressure to give carbamate **5-Et** which was used immediately in the next step without further purification.

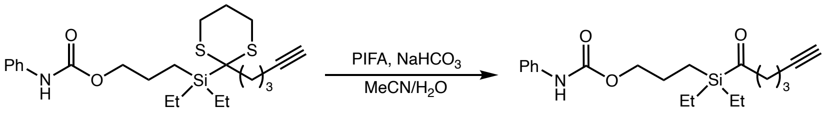


**3-(diethyl(hex-5-ynoyl)silyl)propyl phenylcarbamate (15-Et)**
A scintillation vial was charged with carbamate **5-Et** (0.151 mmol, 1.0 equiv.), $NaHCO_3$ (25 mg, 0.302 mmol, 2.0 equiv.), MeCN (2.72 mL), and water (0.30 mL). PIFA (78 mg, 0.313 mmol, 1.2 equiv.) was added and the reaction was stirred at room temperature. After 30 minutes, the reaction was diluted with $Et_2O$, washed successively with sat. aqueous $NH_4Cl$, water, and brine, and the organic layer was dried over $MgSO_4$, filtered, and concentrated under reduced pressure. The crude material was purified using flash column chromatography (10% EtOAc/Hex) to give acyl silane **15-Et** (5.4 mg, 0.015 mmol, 10% yield) as a clear oil.
**$^{1}$H NMR** (600 MHz, $CDCl_3$) δ 7.42 – 7.35 (m, 2H), 7.34 – 7.27 (m, 2H), 7.06 (tt, *J* = 7.3, 1.2 Hz, 1H), 6.66 (s, 1H), 4.12 (t, *J* = 6.6 Hz, 2H), 2.74 (t, *J* = 7.0 Hz, 2H), 2.20 (td, *J* = 6.9, 2.7 Hz, 2H), 1.95 (t, *J* = 2.6 Hz, 1H), 1.80 – 1.67 (m, 4H), 0.99 (t, *J* = 7.9 Hz, 6H), 0.85 – 0.70 (m, 6H). **$^{13}$C NMR** (151 MHz, $CDCl_3$) δ 247.17, 138.07, 129.20, 129.17, 123.55, 120.79, 118.79, 83.93, 69.13, 48.38, 23.40, 20.64, 17.93, 7.39, 6.63, 2.57. **HRMS** (ESI): calculated for $[C_{20}H_{29}NO_3Si+H]^+$ required *m/z* 360.5485, found *m/z* 360.5421.

**Synthesis of acyl silane 15-cyclo:**

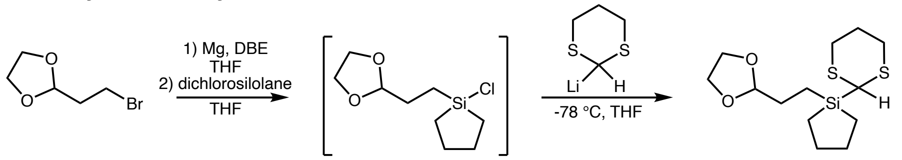


**(2-(1,3-dioxolan-2-yl)ethyl)(1,3-dithian-2-yl)diethylsilane (1-cyclo)**
A flame-dried three-neck flask fitted with a reflux condenser and cooled under $N_2$ was charged with magnesium turnings (1.896 g, 78 mmol, 2.03 equiv.) and THF (60 mL). A small amount of 1,2-dibromoethane was added and the solution was heated with a heat gun until ethylene bubbles were observed. The flask was then placed in a water bath and 2-(2-bromoethyl)-1,3-dioxolane (7.05 mL, 60 mmol, 1.56 equiv.) was added slowly. The mixture was allowed to stir at 40 °C until a majority of the magnesium was consumed (ca. 2 hours). The resulting Grignard reagent was added via cannula to a separate flame-dried flask containing dichlorosilolane (6.05 mL, 46.2 mmol, 1.20 equiv.) and THF (30 mL) at 0 °C. The mixture was allowed to warm to room temperature over 2 hours. Meanwhile, a separate round bottom flask cooled under vacuum was charged with 1,3-dithiane (4.62 g, 38.46 mmol, 1.0 equiv.) and THF (75 mL). The reaction vessel was cooled to –78 ˚C and a solution of *n*BuLi (2.5

M in hexanes, 16.92 mL) was added dropwise resulting in a persistent, but faint yellow color. The flask was allowed to warm to room temperature over 30 minutes and then cooled back to –78 °C and transferred via cannula to the flask containing silyl chloride at –78 °C. The solution was allowed to warm to room temperature overnight. After 16 hours, the reaction was diluted with EtOAc, quenched by addition of sat. aqueous $NH_4Cl$ and washed with water. The combined aqueous layers were then extracted by EtOAc 2X and the combined organic layers were washed with brine, dried over $MgSO_4$, filtered, and concentrated under reduced pressure. The crude oil was then purified via flash-column chromatography (gradient elution 0% – 10% EtOAc/Hex) to give **1-cyclo** as a clear oil (2.879 g, 9.45 mmol, 25% yield).

**$^1$H NMR** (400 MHz, $CDCl_3$) δ 4.85 (q, *J* = 3.5 Hz, 1H), 4.03 – 3.93 (m, 2H), 3.91 – 3.84 (m, 2H), 3.83 (d, *J* = 2.5 Hz, 1H), 2.87 (ddd, *J* = 14.7, 12.3, 2.8 Hz, 2H), 2.70 (dt, *J* = 13.9, 3.6 Hz, 2H), 2.19 – 1.92 (m, 2H), 1.74 (dt, *J* = 12.9, 4.5 Hz, 2H), 1.68 – 1.50 (m, 4H), 0.94 – 0.82 (m, 4H), 0.59 (dt, *J* = 13.0, 6.1 Hz, 2H). **$^{13}$C NMR** (126 MHz, $CDCl_3$) δ 106.06, 65.09, 29.86, 28.06, 26.25, 26.17, 22.83, 12.71, 10.40. **HRMS** (ESI): calculated for $[C_{13}H_{24}O_2S_2Si+Na+MeCN]^+$ required *m/z* 368.1145, found *m/z* 368.1162.

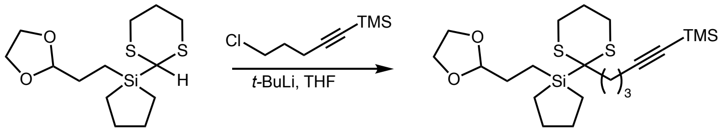


**1-(2-(1,3-dioxolan-2-yl)ethyl)-1-(2-(5-(trimethylsilyl)pent-4-yn-1-yl)-1,3-dithian-2-yl)silolane (2-cyclo)**

A flame-dried round bottom flask cooled under vacuum was charged with **1-cyclo** (2.462 g, 8.20 mmol, 1.0 equiv.) and THF (41 mL, 0.2 M) and cooled to –78 °C, followed by the dropwise addition of *t*BuLi (1.7 M in pentane, 5.01 mL, 1.05 equiv.). After stirring for 30 minutes at –78 °C, the solution was allowed to warm to room temperature over 30 minutes and then cooled back down to –78 °C. Next, (5-chloropent-1-yn-1-yl)trimethylsilane (1.60 mL, 9.02 mmol, 1.1 equiv.) was added dropwise and the reaction was allowed to warm to room temperature overnight. After 16 hours, the reaction was quenched by the addition of sat. aqueous $NH_4Cl$ and extracted with EtOAc. The combined organic layers were washed with saturated brine, dried over $MgSO_4$, and concentrated to an oil. Purification of the crude material by flash column chromatography on silica gel in 95:5 hex/EtOAc yielded the desired product **2-cyclo** as a clear yellow oil (2.169 g, 4.907 mmol, 60% yield).

**$^1$H NMR** (500 MHz, $CDCl_3$) δ 4.82 (t, *J* = 4.6 Hz, 1H), 3.99 – 3.91 (m, 2H), 3.90 – 3.80 (m, 2H), 3.09 (ddd, *J* = 14.9, 12.7, 2.7 Hz, 2H), 2.46 – 2.34 (m, 4H), 2.31 (t, *J* = 6.6 Hz, 2H), 2.09 – 2.00 (m, 1H), 1.91 (qt, *J* = 13.1, 3.4 Hz, 1H), 1.73 – 1.63 (m, 6H), 1.61 – 1.50 (m, 2H), 1.00 (dt, *J* = 14.5, 7.2 Hz, 2H), 0.89 – 0.84 (m, 2H), 0.64 (dt, *J* = 14.3, 6.8 Hz, 2H), 0.16 (s, 9H). **$^{13}$C NMR** (126 MHz, $CDCl_3$) δ 107.24, 105.98, 85.21, 65.09, 38.76, 36.07, 28.74, 27.64, 26.75, 25.29, 23.36, 20.20, 9.84, 5.19, 0.30. **HRMS** (ESI): calculated for $[C_{21}H_{38}O_2S_2Si_2+H]^+$ required *m/z* 443.1925, found *m/z* 443.1992.

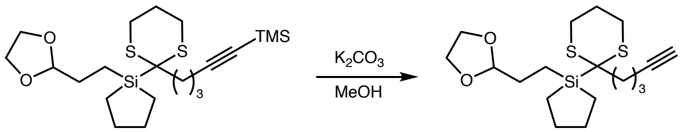


**1-(2-(1,3-dioxolan-2-yl)ethyl)-1-(2-(pent-4-yn-1-yl)-1,3-dithian-2-yl)silolane**
**(3-cyclo)**

A scintillation vial was charged with **2-cyclo** (0.637 g, 1.435 mmol, 1.0 equiv.), potassium carbonate (0.595 g, 4.30 mmol, 3 equiv.), and MeOH (3.59 mL). The reaction was allowed to stir at room temperature for 90 minute and was then diluted with ether and washed with water and brine. The organic layers were dried over $MgSO_4$, filtered, and concentrated under reduced pressure to give alkyne **3-cyclo** (0.452 g, 1.215 mmol, 85% yield) as a clear oil.

**$^1$H NMR** (500 MHz, $CDCl_3$) δ 4.83 (t, *J* = 4.6 Hz, 1H), 4.02 – 3.91 (m, 2H), 3.91 – 3.80 (m, 2H), 3.07 (ddd, *J* = 15.0, 12.6, 2.7 Hz, 2H), 2.42 (ddd, *J* = 14.2, 4.2, 3.1 Hz, 2H), 2.39 – 2.34 (m, 2H), 2.26 (td, *J* = 6.8, 2.6 Hz, 2H), 2.09 – 2.02 (m, 1H), 1.99 (t, *J* = 2.6 Hz, 1H), 1.90 (qt, *J* = 13.0, 3.3 Hz, 1H), 1.74 – 1.63 (m, 6H), 1.57 – 1.50 (m, 3H), 1.00 (dt, *J* = 14.6, 7.4 Hz, 2H), 0.90 – 0.85 (m, 2H), 0.64 (dt, *J* = 14.3, 6.9 Hz, 2H). **$^{13}$C NMR** (126 MHz,

$CDCl_3$) δ 105.98, 84.33, 68.98, 65.12, 38.68, 36.27, 28.75, 27.63, 26.79, 25.28, 23.43, 18.82, 9.94, 5.16. **HRMS** (ESI): calculated for $[C_{18}H_{30}O_2S_2Si+H]^+$ required *m/z* 371.1529, found *m/z* 371.1532.

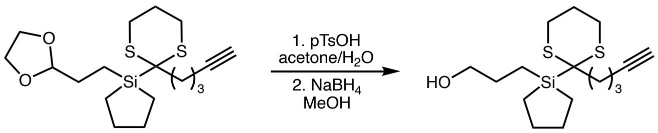


**3-(1-(2-(pent-4-yn-1-yl)-1,3-dithian-2-yl)silolan-1-yl)propan-1-ol (4-cyclo)**
A 50 mL round-bottom flask was charged with alkyne **3-cyclo** (0.7862 g, 2.121 mmol, 1.0 equiv.) and acetone (28.3 mL). Water (7.1 mL) and *p*-TsOH (2.191 g, 12.726 mmol, 6.0 equiv.) were added and the reaction was stirred at room temperature. After 16 hours, the reaction was quenched by addition of sat. aqueous $NaHCO_3$ and extracted with EtOAc. The combined organic layers were washed with brine, dried over $MgSO_4$, filtered, and concentrated to give the aldehyde was used immediately in the next step without further purification. A scintillation vial was then charged with aldehyde and MeOH (8.5 mL). The vial was cooled to 0 °C and sodium borohydride (67 mg) was added and the reaction was allowed to stir. After one hour, an additional dose of sodium borohydride (40 mg) was added and the reaction was stirred for an additional hour. The reaction was then quenched by addition of water and extracted with EtOAc. The combined organic layers were washed with brine, dried over $MgSO_4$, filtered, and concentrated under reduced pressure. The crude oil was purified using flash column chromatography (25% EtOAc/hex) to give alcohol **4-cyclo** (0.380 g, 1.156 mmol, 55% yield over two steps) as a clear oil.
**$^1$H NMR** (500 MHz, $CDCl_3$): δ 3.62 (t, *J* = 6.6 Hz, 2H), 3.07 (ddd, *J* = 15.0, 12.7, 2.7 Hz, 2H), 2.43 (ddd, *J* = 14.3, 4.2, 3.1 Hz, 2H), 2.40 – 2.33 (m, 2H), 2.27 (td, *J* = 6.7, 2.7 Hz, 2H), 2.10 – 2.03 (m, 1H), 1.99 (t, *J* = 2.7 Hz, 1H), 1.90 (qt, *J* = 12.9, 3.3 Hz, 1H), 1.73 – 1.65 (m, 4H), 1.66 – 1.60 (m, 2H), 1.59 – 1.52 (m, 2H), 1.02 (dt, *J* = 14.6, 7.2 Hz, 2H), 0.83 – 0.75 (m, 2H), 0.64 (dt, *J* = 14.5, 6.9 Hz, 2H). **$^{13}$C NMR** (126 MHz, $CDCl_3$): δ 84.33, 68.98, 65.74, 38.71, 36.33, 27.66, 27.58, 26.78, 25.28, 23.42, 18.82, 10.03, 7.23. **HRMS** (ESI): calculated for $[C_{16}H_{28}OS_2Si+H]^+$ required *m/z* 329.1424, found *m/z* 329.1450.

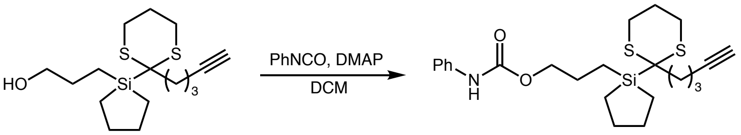


**3-(1-(2-(pent-4-yn-1-yl)-1,3-dithian-2-yl)silolan-1-yl)propyl phenylcarbamate (5-cyclo)**
A scintillation vial was charged with alcohol **4-cyclo** (0.150 g, 0.456 mmol, 1.0 equiv.), DMAP (6 mg) and DCM (2.28 mL). Phenylisocyanate (55 uL, 0.502 mmol, 1.1 equiv.) was added and the reaction was allowed to stir at room temperature overnight. After 16 hours, the reaction was diluted with DCM, washed successively with sat. aqueous $NH_4Cl$, water, and brine, and the organic layer was dried over $MgSO_4$, filtered, and concentrated under reduced pressure to give carbamate **5-cyclo** (99% yield) which was used immediately in the next step without further purification.

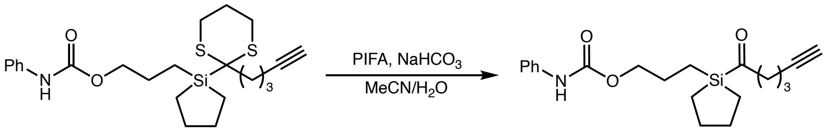


**3-(1-(hex-5-ynoyl)silolan-1-yl)propyl phenylcarbamate (15-cyclo)**
A scintillation vial was charged with carbamate **5-cyclo** (0.456 mmol, 1.0 equiv.), $NaHCO_3$ (77 mg, 0.912 mmol, 2.0 equiv.), MeCN (8.21 mL), and water (0.91 mL). PIFA (0.235 g, 0.547 mmol, 1.2 equiv.) was added and the reaction was stirred at room temperature. After 30 minutes, the reaction was diluted with $Et_2O$, washed successively with sat. aqueous $NH_4Cl$, water, and brine, and the organic layer was dried over $MgSO_4$, filtered,

and concentrated under reduced pressure. The crude material was purified using preparatory TLC (10% EtOAc/Hex) to give acyl silane **15-cyclo** (32 mg, 0.090 mmol, 20% yield) as a white solid.

**$^1$H NMR** (400 MHz, MeOD) δ 7.41 (d, *J* = 8.0 Hz, 2H), 7.31 – 7.21 (m, 2H), 7.06 – 6.96 (m, 1H), 4.09 (t, *J* = 6.6 Hz, 2H), 2.82 (t, *J* = 7.0 Hz, 2H), 2.22 (q, *J* = 2.8 Hz, 1H), 2.15 (td, *J* = 7.0, 2.7 Hz, 2H), 1.80 – 1.59 (m, 8H), 0.98 – 0.81 (m, 4H), 0.80 – 0.68 (m, 2H). **$^{13}$C NMR** (151 MHz, MeOD) δ 248.60, 156.05, 140.17, 129.79, 124.00, 119.87, 84.40, 70.22, 67.98, 28.32, 24.72, 22.10, 18.40, 10.47, 8.52. **HRMS** (ESI): calculated for $[C_{20}H_{27}NO_3Si+H]^+$ required *m/z* 358.5325, found *m/z* 358.5286.

**Synthesis of acyl silane JQ1-Et**

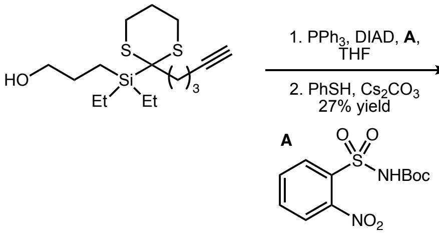


***tert*-butyl (3-(diethyl(2-(pent-4-yn-1-yl)-1,3-dithian-2-yl)silyl)propyl)carbamate (6-Et)**

A scintillation vial equipped with a magnetic stir bar was charged with alcohol **4-Et** (0.086 g, 0.260 mmol, 1.0 equiv.), reagent **A** (0.090 g, 0.299 mmol, 1.15 equiv.), triphenylphosphine (0.079 g, 0.299 mmol, 1.15 equiv.) and THF (1.3 mL). A solution of DIAD (59 µL, 0.299 mmol, 1.15 equiv.) in THF (1.3 mL) was added at 0 °C and the mixture was allowed to stir at room temperature. After 36 hours, cesium carbonate (0.320 g, 0.983 mmol, 3.75 equiv.) and thiophenol (55 µL, 0.546 mmol, 2.1 equiv.) were added and the reaction was heated to 50 °C. After an additional 4 hours, the reaction was quenched by addition of 0.5 M NaOH and extracted thrice with EtOAc. The combined organic layers were washed with brine, dried over $MgSO_4$, filtered, and concentrated *in vacuo*. The residue was purified by preparatory TLC (10% MeOH/DCM) to give *N*-boc protected amine **6-Et** (0.030 g, 0.070 mmol, 27% yield) which was used immediately in the next step without further purification.

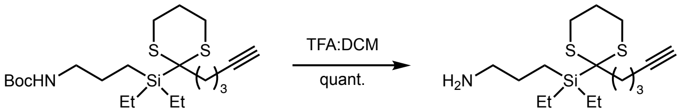


**3-(diethyl(2-(pent-4-yn-1-yl)-1,3-dithian-2-yl)silyl)propan-1-amine (7-Et)**

A scintillation vial was charged with amine **6-Et** (0.030 g, 0.070 mmol) and DCM (1 mL). The mixture was cooled to 0 °C at which point trifluoroacetic acid (83 µL) was added and the reaction was allowed to stir at room temperature for 1 hour. The mixture was concentrated *in vacuo* to give amine **7-Et** (0.026 g, 0.070 mmol, 99% yield) as an oil.

**$^1$H NMR** (500 MHz, $CDCl_3$) δ 3.07 (ddd, *J* = 15.1, 12.8, 2.7 Hz, 2H), 2.76 (t, *J* = 7.1 Hz, 2H), 2.38 (ddt, *J* = 12.4, 8.6, 4.0 Hz, 4H), 2.27 (td, *J* = 6.7, 2.7 Hz, 2H), 2.06 – 1.98 (m, 1H) 1.99 (t, *J* = 2.6 Hz, 1H), 1.97 – 1.85 (m, 1H), 1.77 – 1.63 (m, 4H), 1.04 (t, *J* = 7.9 Hz, 2H), 0.82 – 0.76 (m, 6H). **$^{13}$C NMR** (126 MHz, $CDCl_3$) δ 84.29, 68.93, 44.91, 39.39, 36.61, 26.87, 26.81, 25.19, 23.48, 18.84, 8.21, 8.16, 3.26, 1.10 *(some impurity observed)*. **HRMS** (ESI): calculated for $[C_{16}H_{31}NS_2Si+H]^+$ required *m/z* 330.1740, found *m/z* 330.1745.

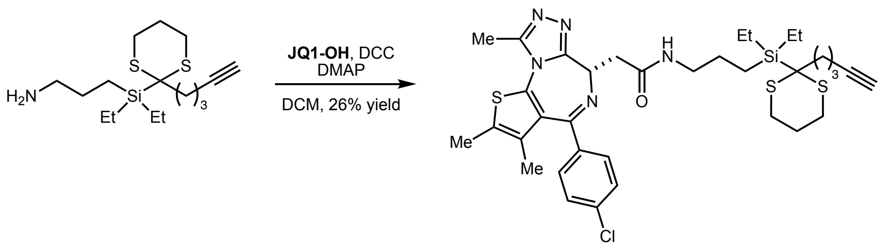


**(*S*)-2-(4-(4-chlorophenyl)-2,3,9-trimethyl-6*H*-thieno[3,2-*f*][1,2,4]triazolo[4,3-*a*][1,4]diazepin-6-yl)-*N*-(3-(diethyl(2-(pent-4-yn-1-yl)-1,3-dithian-2-yl)silyl)propyl)acetamide (8-Et)**

A one-dram vial was charged with **JQ1-OH** (10 mg, 0.025 mmol, 1.1 equiv.), dicyclohexyl carbodiimide (5.4 mg, 0.026 mmol, 1.15 equiv.), and DMAP (5.4 mg, 0.011 mmol, 0.5 equiv.). The reaction vessel was evacuated and backfilled with nitrogen three times after which a solution of amine **7-Et** (7.5 mg, 0.023 mmol, 1.0 equiv.) in DCM (0.23 mL) was added. The mixture was allowed to stir at room temperature. After 16 hours, the mixture was concentrated *in vacuo* and purified using preparatory TLC (5% MeOH/DCM) to give dithiane **8-Et** (4 mg, 0.006 mmol, 26% yield) as a white solid.

**$^1$H NMR** (600 MHz, $CDCl_3$) δ 7.40 (d, *J* = 8.2 Hz, 2H), 7.33 (d, *J* = 8.2 Hz, 2H), 6.51 – 6.46 (m, 1H), 4.63 (t, *J* = 7.0 Hz, 1H), 3.55 (dd, *J* = 14.2, 7.2 Hz, 1H), 3.49 (s, 1H), 3.35 (ddd, *J* = 14.2, 6.6, 4.5 Hz, 2H), 3.24 (dq, *J* = 13.0, 6.6 Hz, 1H), 3.12 – 3.04 (m, 2H), 2.67 (s, 3H), 2.44 – 2.35 (m, 4H), 2.40 (s, 3H), 2.28 (td, *J* = 6.7, 2.6 Hz, 2H), 2.08 – 2.01 (m, 1H), 2.03 (t, *J* = 2.6 Hz, 1H), 1.99 – 1.88 (m, 2H), 1.77 – 1.67 (m, 4H), 1.67 (s, 3H), 1.40 – 1.30 (m, H), 1.20 – 1.06 (m, 4H), 1.04 (td, *J* = 7.9, 3.2 Hz, 4H), 0.84 – 0.76 (m, 6H), 0.73 – 0.65 (m, 1H). **$^{13}$C NMR** (151 MHz, $CDCl_3$) δ 170.49, 164.02, 136.93, 136.82, 129.97, 128.89, 84.37, 69.08, 54.69, 43.16, 39.70, 39.50, 36.71, 34.10, 26.89, 25.77, 25.28, 25.09, 24.61, 23.57, 18.92, 14.52, 13.23, 11.98, 8.53, 8.31, 3.35, 3.33 *(one quaternary resonance not observed)*. **HRMS** (ESI): calculated for $[C_{35}H_{46}ClN_5OS_3Si+H]^+$ required *m/z* 712.2395, found *m/z* 712.2413.

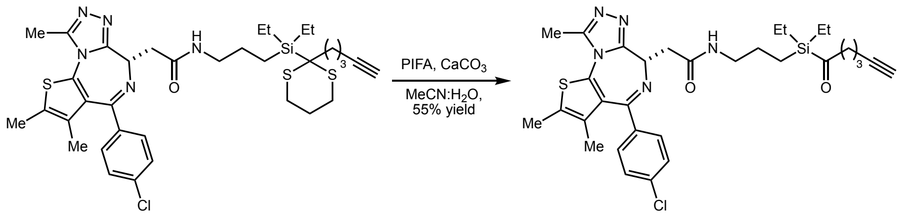


**(*S*)-2-(4-(4-chlorophenyl)-2,3,9-trimethyl-6*H*-thieno[3,2-*f*][1,2,4]triazolo[4,3-*a*][1,4]diazepin-6-yl)-*N*-(3-(diethyl(hex-5-ynoyl)silyl)propyl)acetamide (JQ1-Et)**

A one-dram vial was charged with dithiane **8-Et** (6.62 mg, 9.29 µmol, 1.0 equiv.), PIFA (4.63 mg, 9.76 µmol, 1.05 equiv.), and $CaCO_3$ (9.29 mg, 92.90 µmol, 10 equiv.). A solution of MeCN/$H_2O$ (306 µL/34 µL) was added and the reaction was allowed to stir at room temperature for 3 hours. The mixture was quenched by addition of sat. aq. $NaHCO_3$ and extracted thrice with ethyl acetate. The combined organic layers were dried over $MgSO_4$, filtered, and concentrated *in vacuo*. The crude residue was purified using preparatory TLC to give acyl silane **JQ1-Et** (3.18 mg, 5.11 µmol, 55% yield) as a white solid.

**$^1$H NMR** (600 MHz, $CDCl_3$) δ 7.40 (d, *J* = 8.3 Hz, 2H), 7.33 (d, *J* = 8.4 Hz, 3H), 6.61 (s, 1H), 4.61 (dd, *J* = 7.8, 6.1 Hz, 1H), 3.57 (dd, *J* = 14.1, 7.8 Hz, 1H), 3.38 – 3.16 (m, 3H), 2.72 (t, *J* = 7.0 Hz, 2H), 2.68 (s, 3H), 2.40 (s, 3H), 2.19 (td, *J* = 6.9, 2.7 Hz, 2H), 1.97 (t, *J* = 2.6 Hz, 1H), 1.73 (p, *J* = 7.0 Hz, 2H), 1.67 (s, 3H), 1.57 – 1.51 (m, 2H), 0.96 (td, *J* = 7.9, 4.1 Hz, 6H), 0.77 – 0.69 (m, 6H). **$^{13}$C NMR** (151 MHz, $CDCl_3$) δ 246.96, 170.51, 164.05, 155.75, 136.98, 136.68, 131.05, 129.93, 128.87, 83.93, 69.11, 54.66, 48.43, 42.70, 39.67, 32.03, 23.83, 22.80, 20.62, 17.89, 14.47, 14.22, 13.20, 11.93, 7.53, 7.39, 2.54. **HRMS** (ESI): calculated for $[C_{32}H_{41}ClN_5O_2SSi+H]^+$ required *m/z* 622.2433, found *m/z* 622.2442.

### NMR spectra

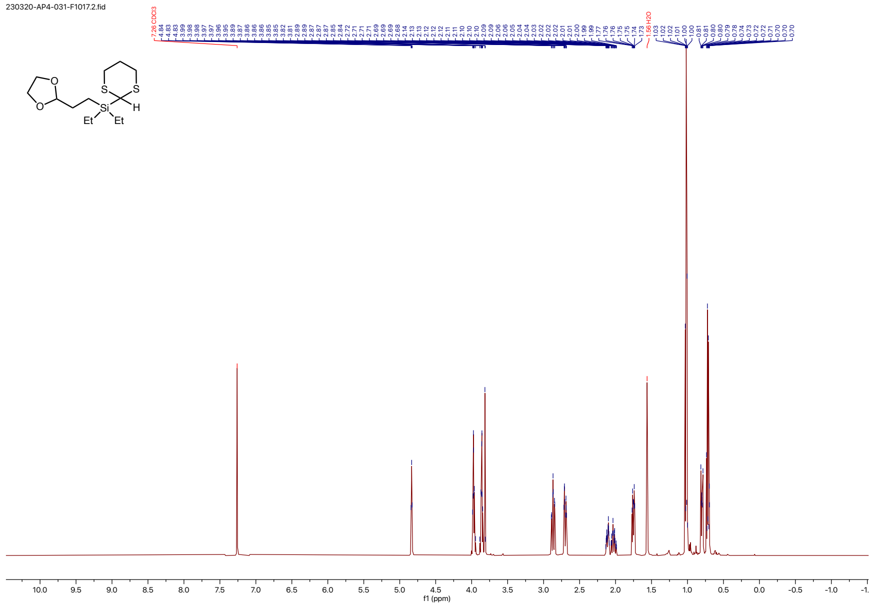

230320-AP4-031-F1017.3.fid

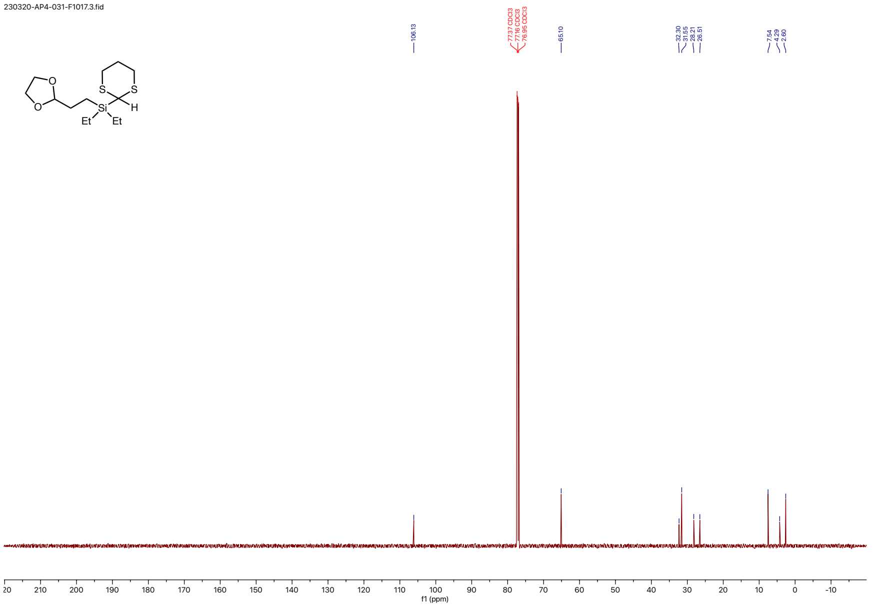

230323-AP4-032B-F2023.2.fid

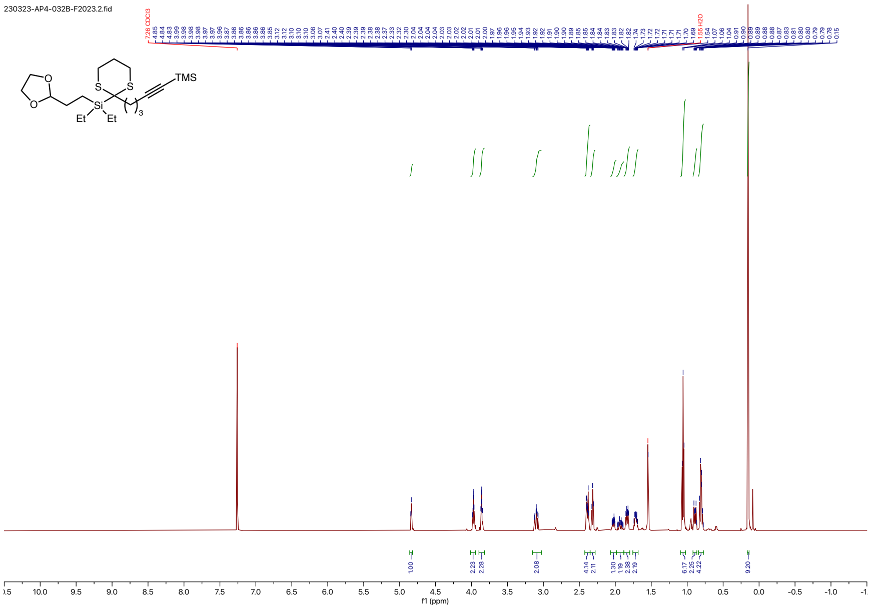

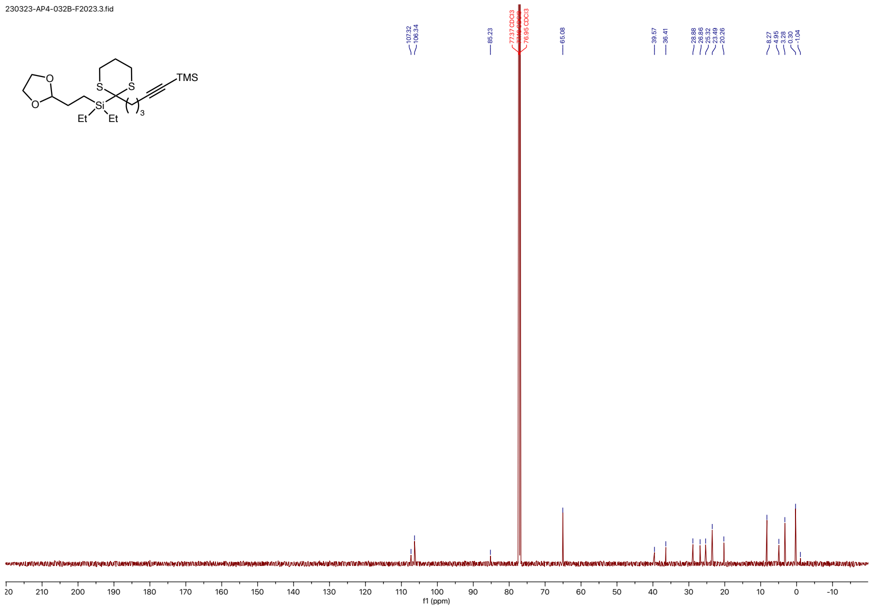
230323-AP4-032B-F2023.3.fid
TMS
S
S
O
O
Si
Et
Et
3
107.32
106.34
85.23
77.37 CDCl3
76.95 CDCl3
65.08
39.57
36.41
28.88
26.86
25.32
23.49
20.26
8.27
4.95
3.28
0.30
-1.04
20
210
200
190
180
170
160
150
140
130
120
110
100
90
80
70
60
50
40
30
20
10
0
-10
f1 (ppm)

240729-AP5-062-F1018.10.fid

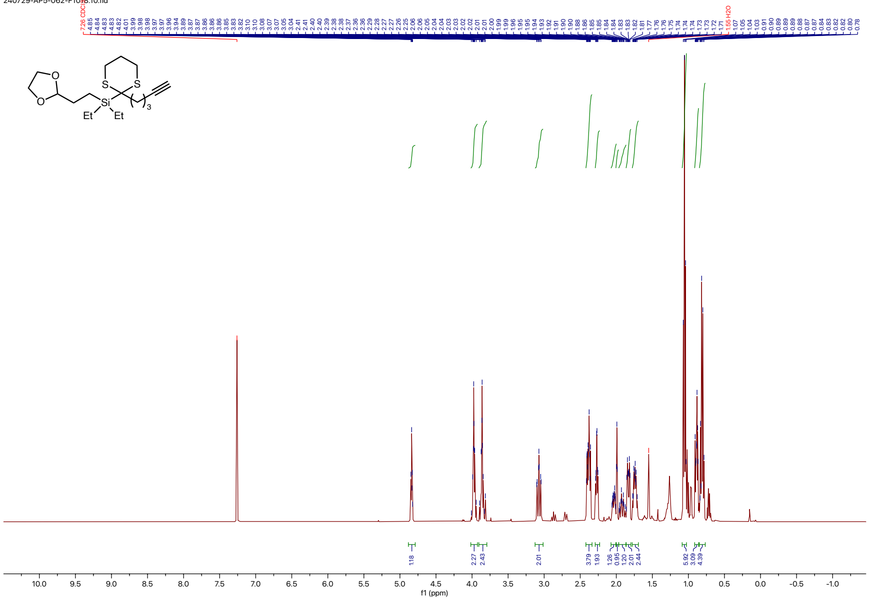

240801-AP5-062-p.10.fid

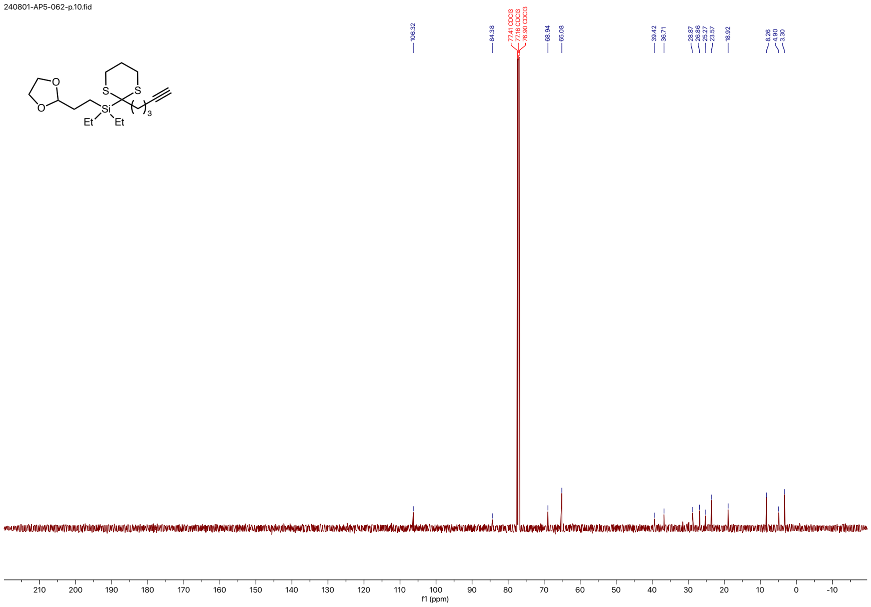

240731-AP5-066-F1317.10.fid

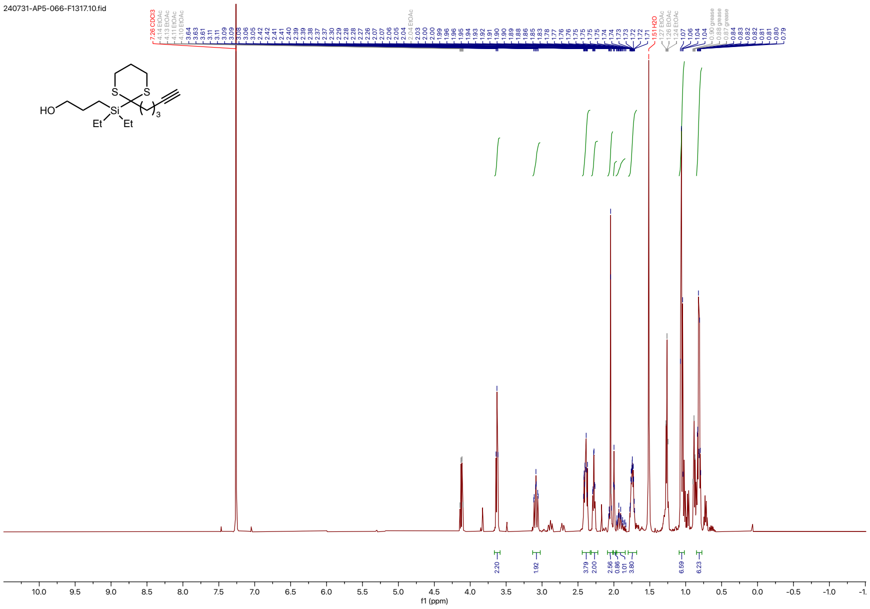

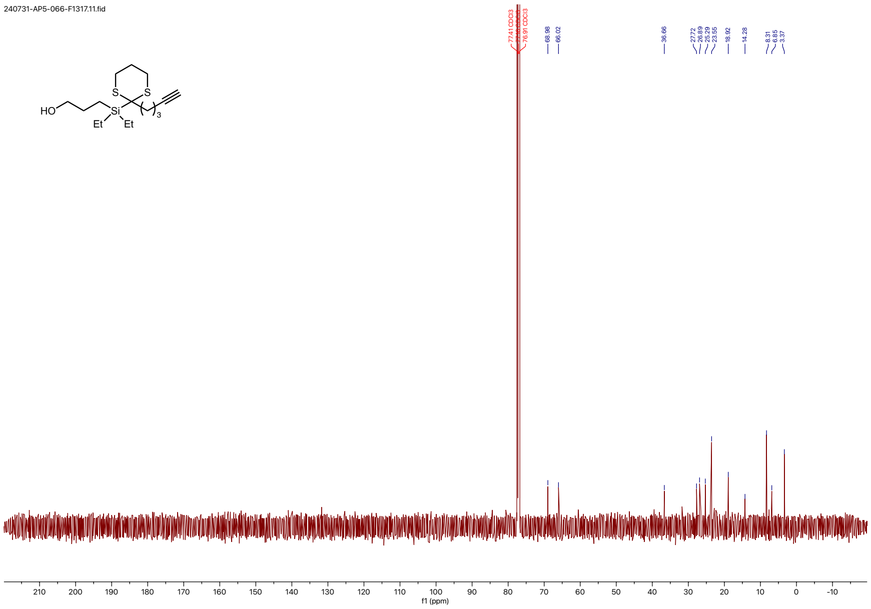

240731-AP5-066-F1317.11.fid
77.41 CDCl3
76.91 CDCl3
68.98
66.02
36.66
27.72
26.89
25.29
23.55
18.92
14.28
8.31
6.85
3.37
HO
S
S
Si
Et
Et
3
210
200
190
180
170
160
150
140
130
120
110
100
90
80
70
60
50
40
30
20
10
0
-10
f1 (ppm)

240802-AP5-072-F13-xx_oil_cdcl3.2.fid

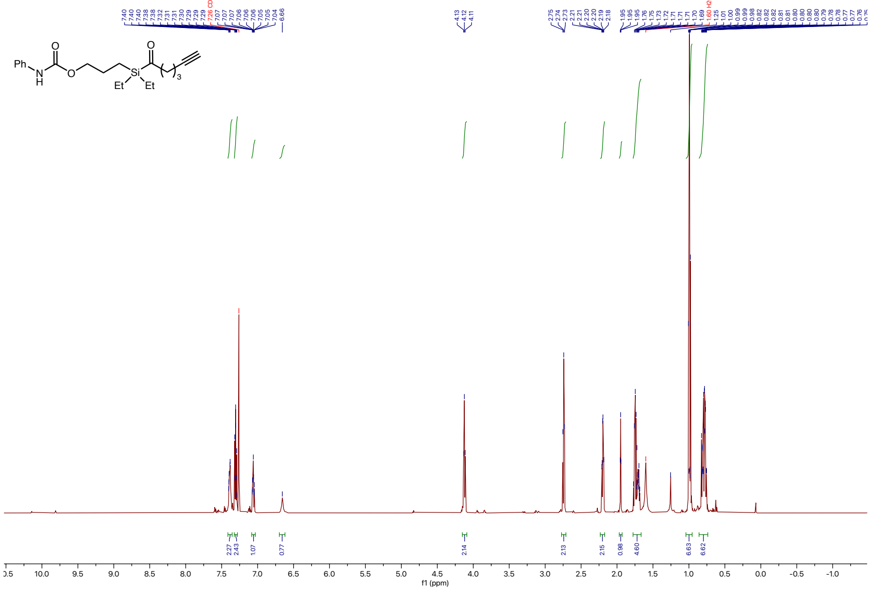

240802-AP5-073-carbon.1.fid

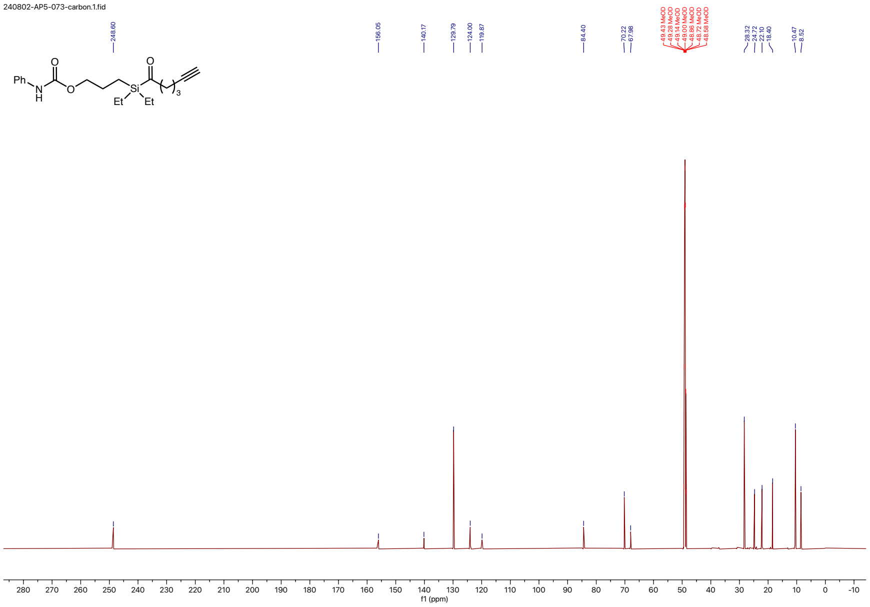

230709-AP4-123-F5370.1.fid

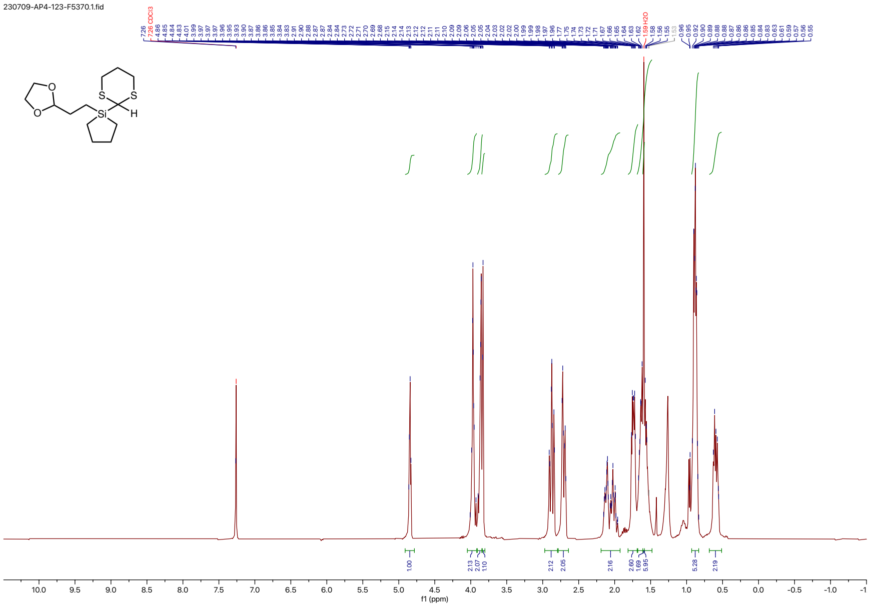

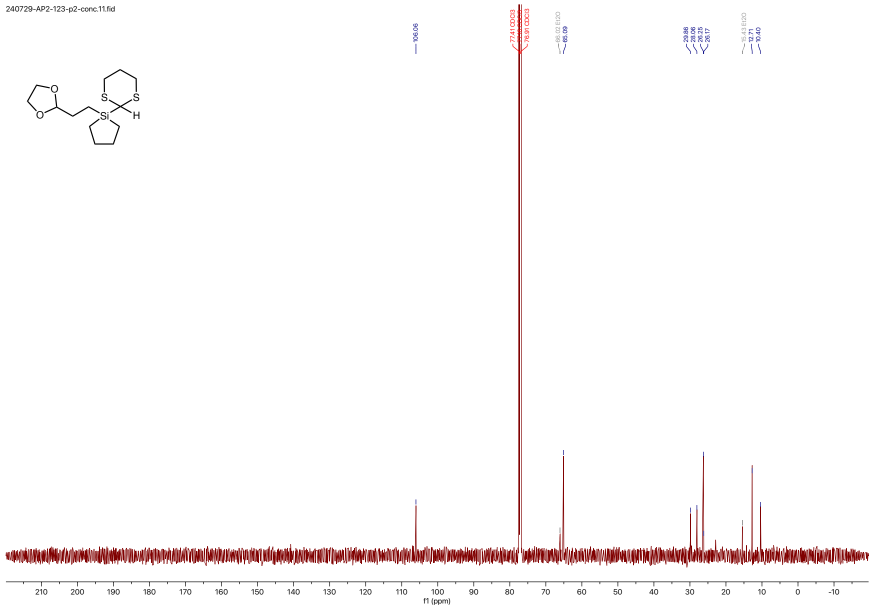

240729-AP2-123-p2-conc.11.fid
106.06
77.41 CDCl3
76.91 CDCl3
65.88 Et2O
65.09
29.86
28.06
26.25
26.17
15.42 Et2O
12.71
10.40
O
O
S
S
Si
H
210
200
190
180
170
160
150
140
130
120
110
100
90
80
70
60
50
40
30
20
10
0
-10
f1 (ppm)

240728-AP2-125-p.10.fid

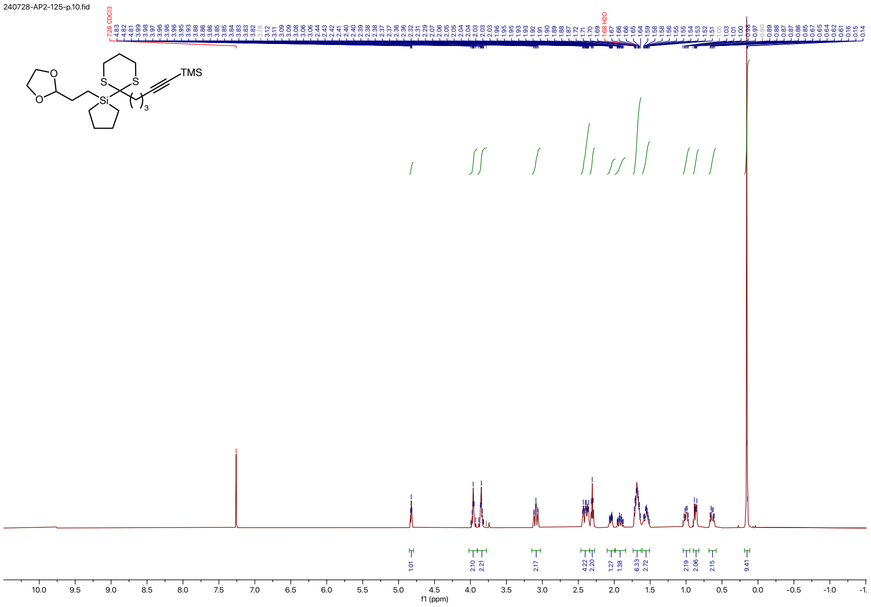

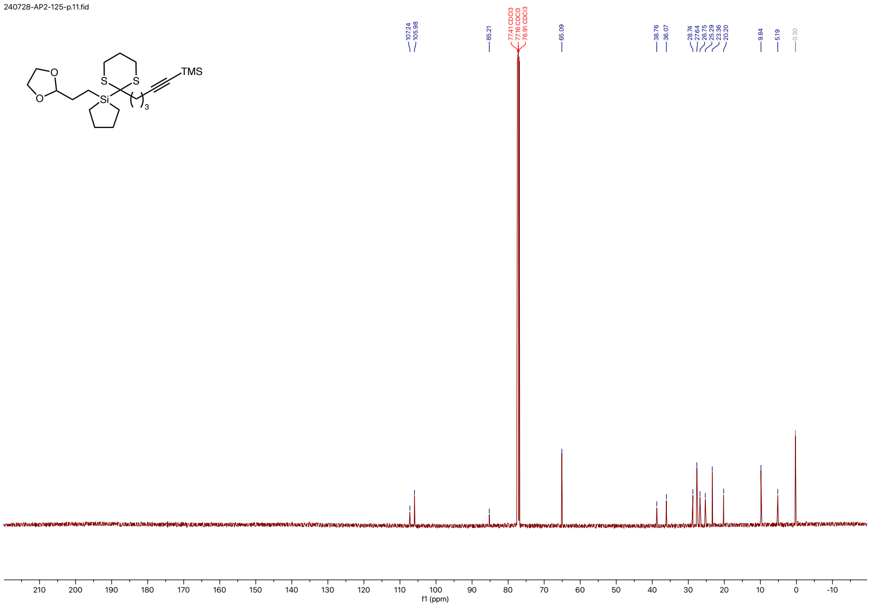
240728-AP2-125-p.11.fid
O
O
S
S
Si
TMS
3
107.24
105.98
85.21
77.41 CDCl3
77.16 CDCl3
76.91 CDCl3
65.09
38.76
36.07
28.74
27.64
26.75
25.29
23.36
20.20
9.84
5.19
0.30
210
200
190
180
170
160
150
140
130
120
110
100
90
80
70
60
50
40
30
20
10
0
-10
f1 (ppm)

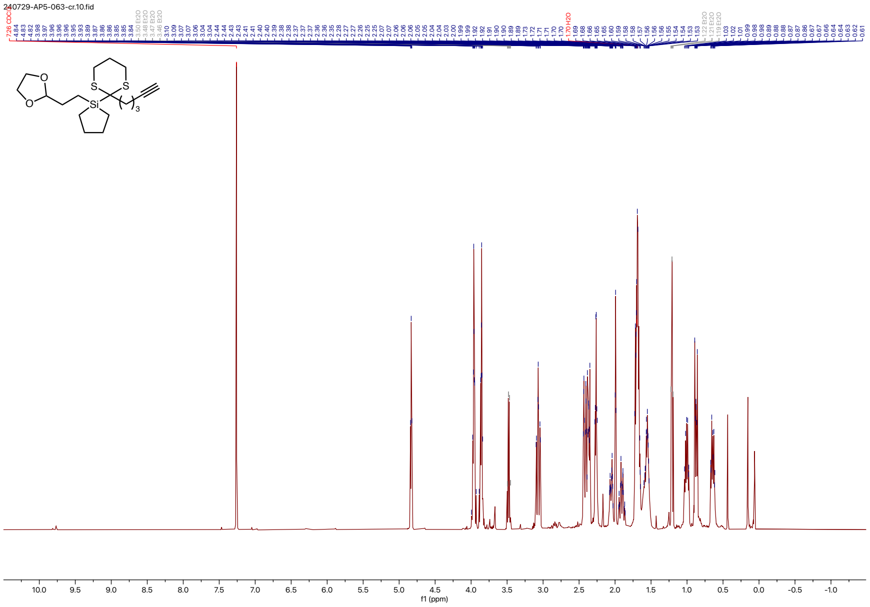
240729-AP5-063-cr.10.fid
10.0 9.5 9.0 8.5 8.0 7.5 7.0 6.5 6.0 5.5 5.0 4.5 4.0 3.5 3.0 2.5 2.0 1.5 1.0 0.5 0.0 -0.5 -1.0
f1 (ppm)

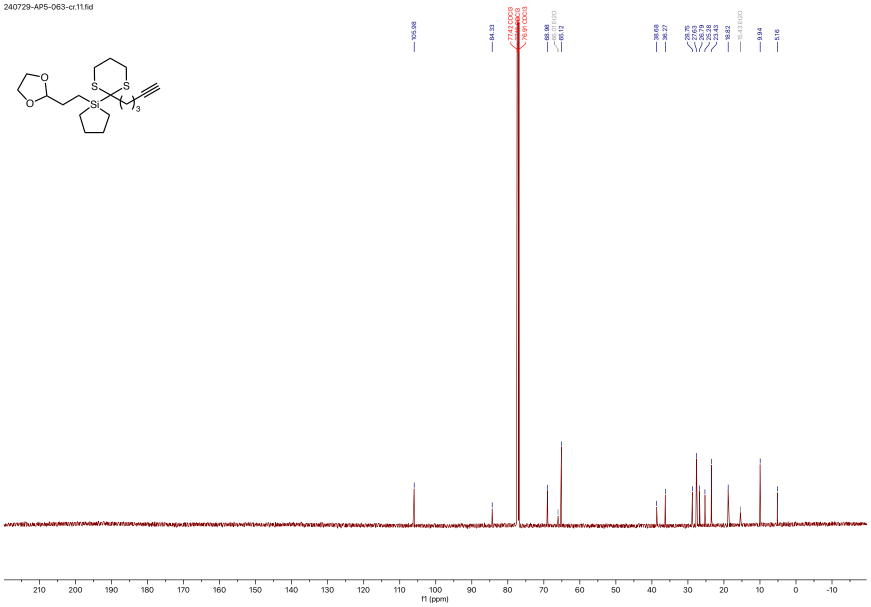

240729-AP5-063-cr.11.fid
O
O
S
S
Si
3
105.98
84.33
77.42 CDCl3
77.16 CDCl3
76.91 CDCl3
68.98
66.01 Et2O
65.12
38.68
36.27
28.75
27.63
26.79
25.28
23.43
18.82
15.43 Et2O
9.94
5.16
210 200 190 180 170 160 150 140 130 120 110 100 90 80 70 60 50 40 30 20 10 0 -10
f1 (ppm)

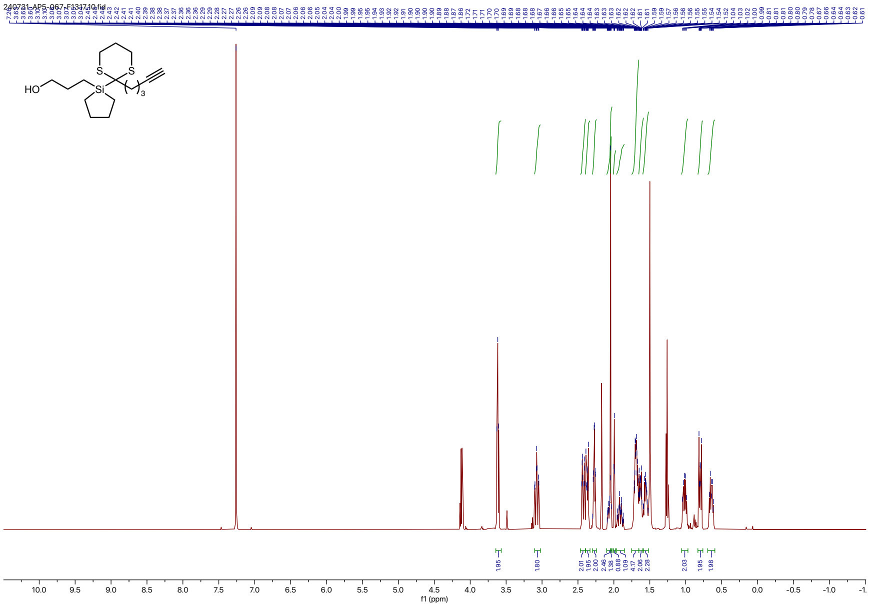
240731-AP5-067-F131710.fid
HO
S
S
Si
3
f1 (ppm)

240731-AP5-067-F1317.11.fid

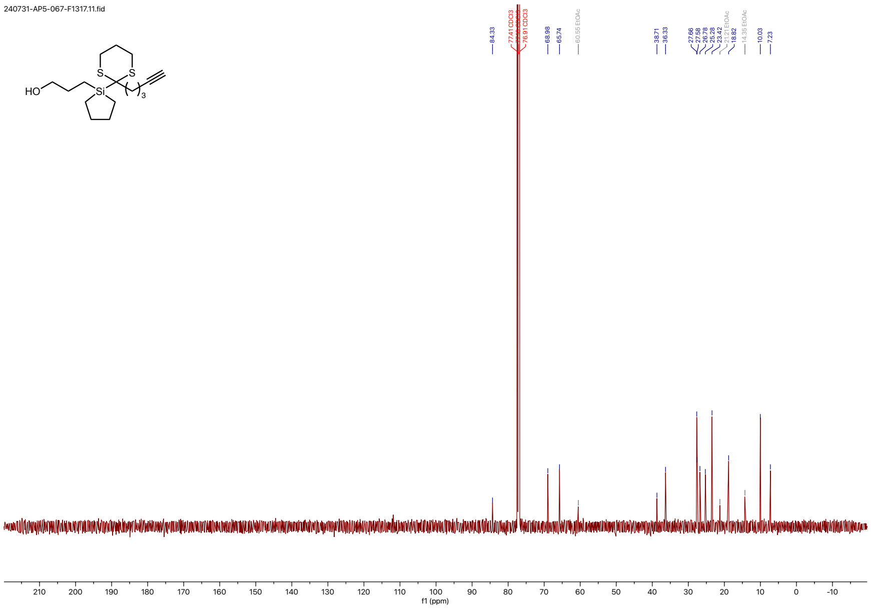

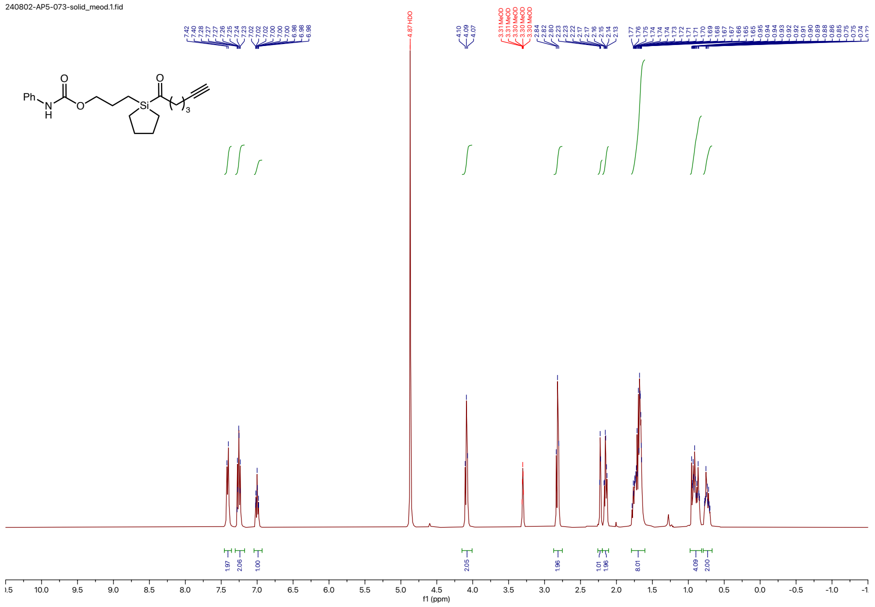
240802-AP5-073-solid_meod.1.fid
f1 (ppm)

240802-AP5-073-carbon.1.fid

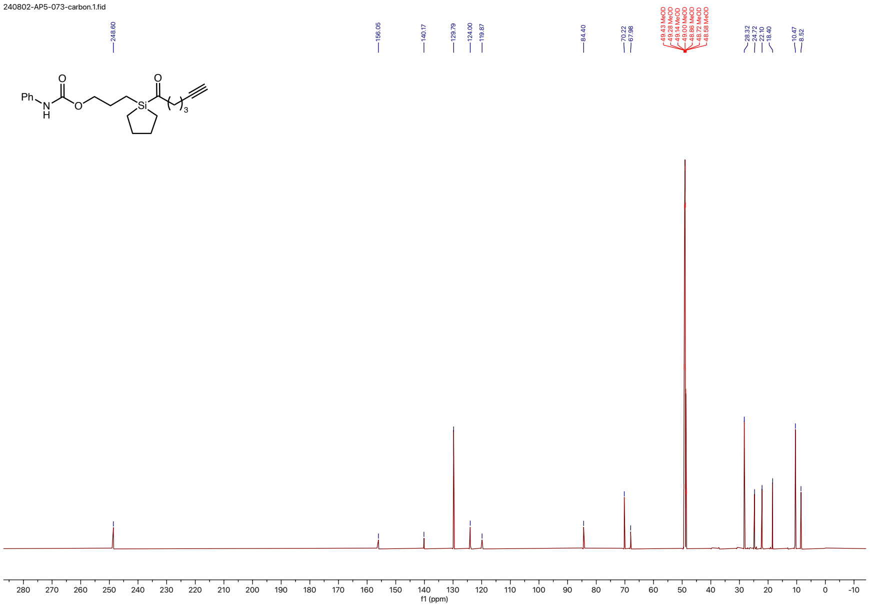

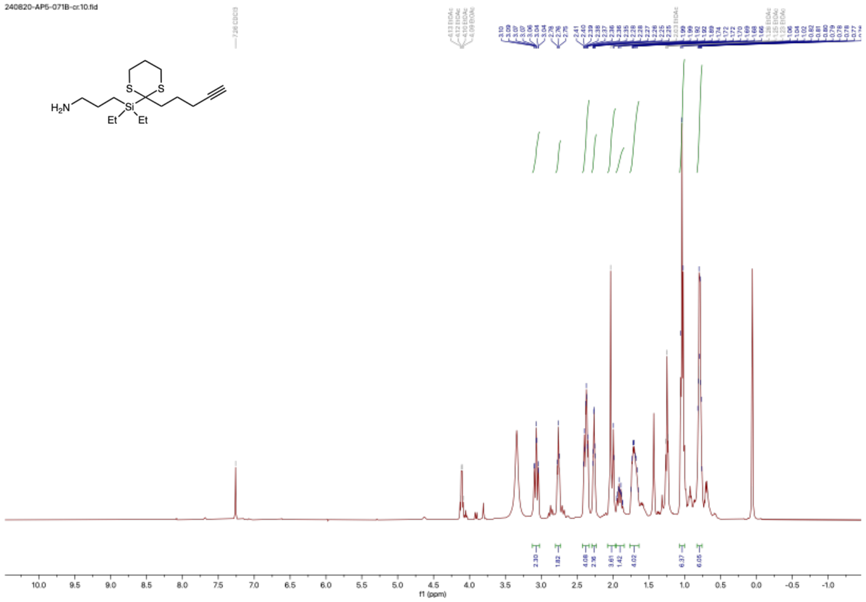
240820-AP5-071B-cr.10.fid
H2N
S
S
Si
Et
Et
2.30
1.82
4.08
2.36
3.61
1.42
4.02
6.37
6.05
10.0
9.5
9.0
8.5
8.0
7.5
7.0
6.5
6.0
5.5
5.0
4.5
4.0
3.5
3.0
2.5
2.0
1.5
1.0
0.5
0.0
-0.5
-1.0
f1 (ppm)

240820-AP5-071B-cr.11.fid

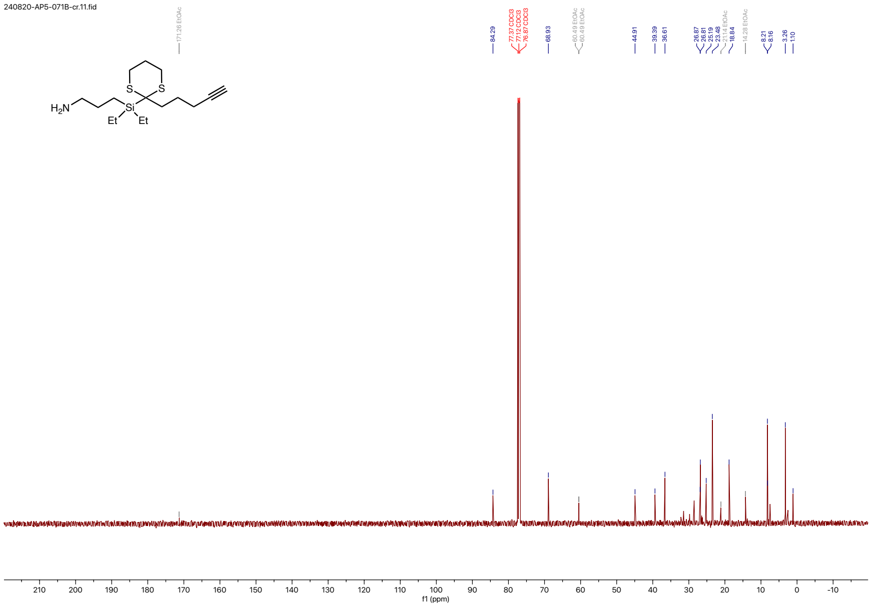

240822-AP5-075-P.4.fid

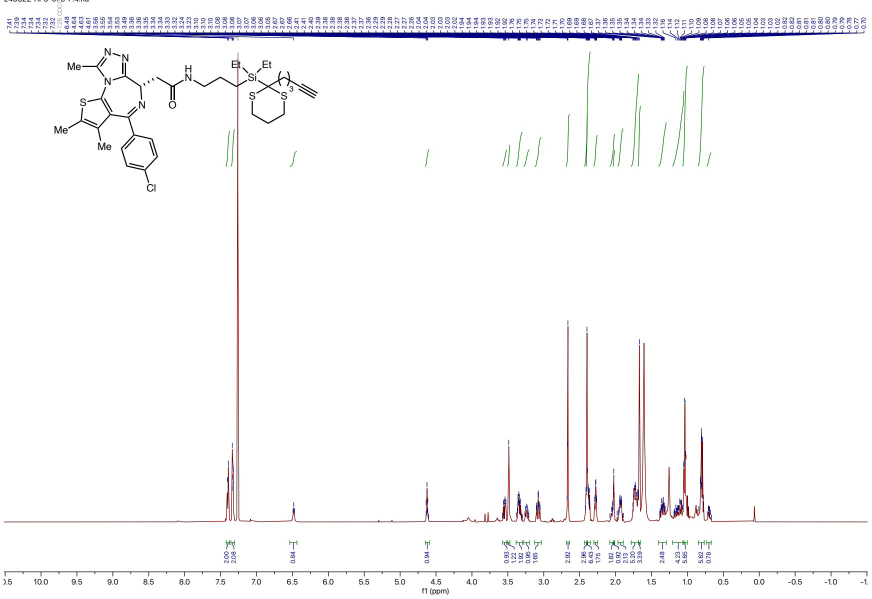

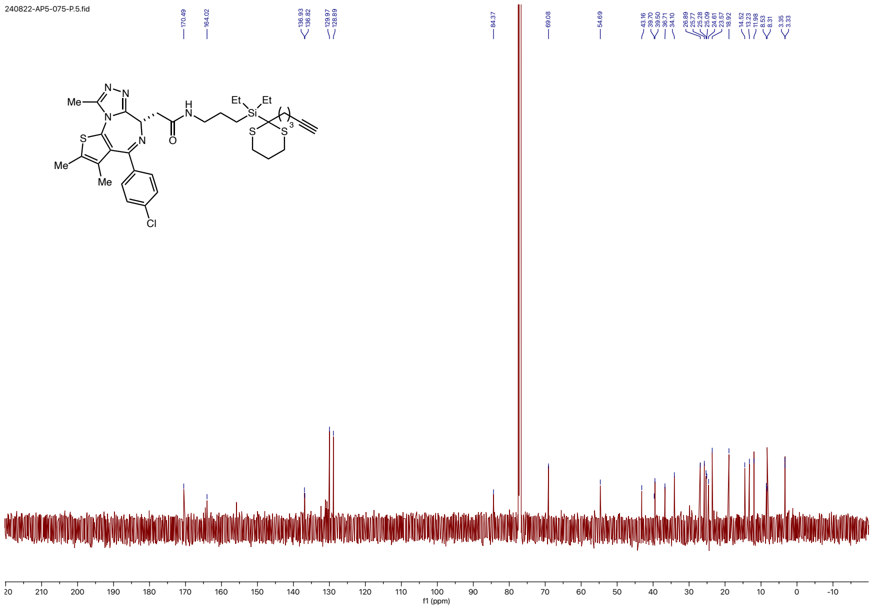
240822-AP5-075-P.5.fid
170.49
164.02
136.93
136.82
129.97
128.89
84.37
69.08
54.69
43.16
39.70
39.50
36.71
34.10
26.89
25.77
25.28
25.09
24.61
23.57
18.92
14.52
13.23
11.98
8.53
8.31
3.35
3.33
Me
N
N
H
N
Et
Et
Si
3
S
S
N
O
S
N
Me
Me
Cl
210
200
190
180
170
160
150
140
130
120
110
100
90
80
70
60
50
40
30
20
10
0
-10
f1 (ppm)

240908-AP5-jq1_et_carbon.4.fid
1H starting parameters - HC 06/17/2019
No decoupling

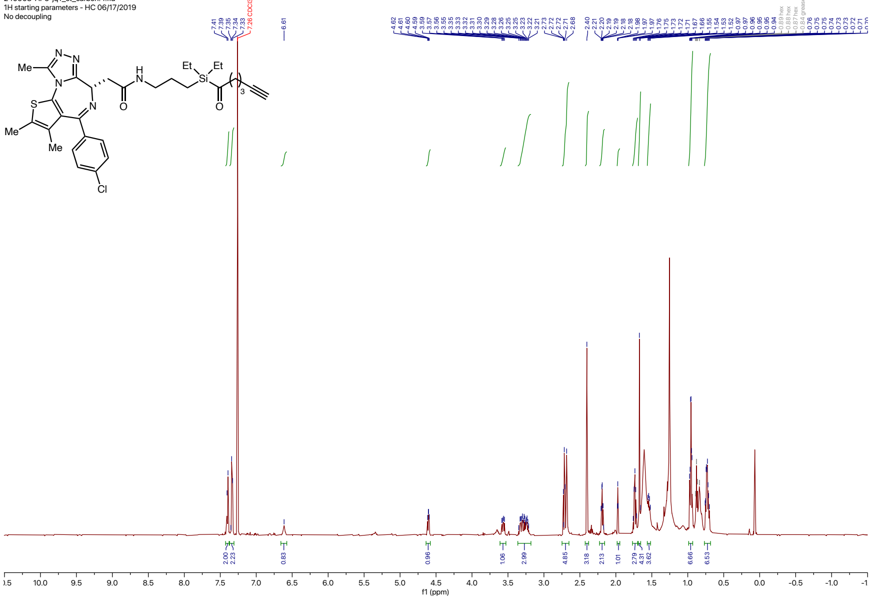

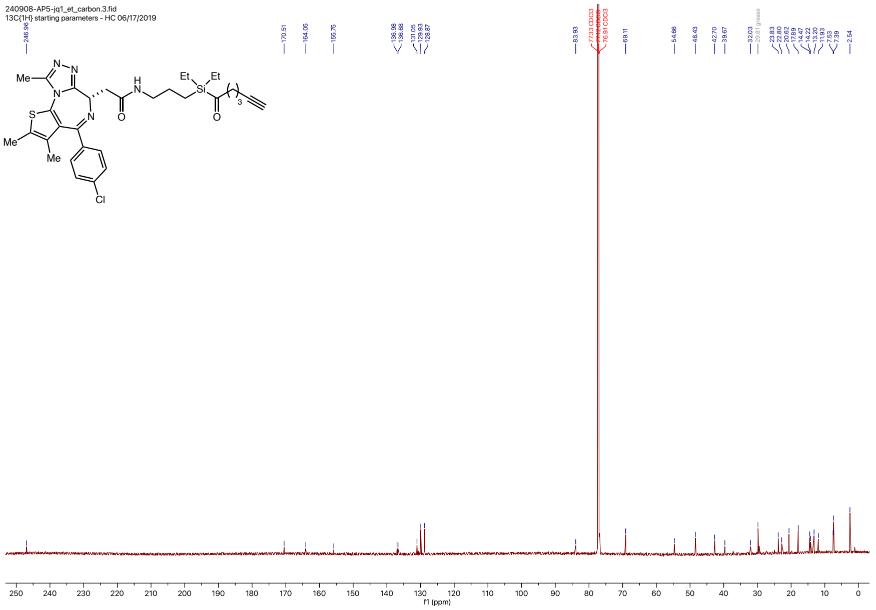
240908-AP5-jq1_et_carbon.3.fid
13C{1H} starting parameters - HC 06/17/2019
246.96
170.51
164.05
155.75
136.98
136.68
131.05
129.93
128.87
83.93
77.33 CDCl3
77.12 CDCl3
76.91 CDCl3
69.11
54.66
48.43
42.70
39.67
32.03
29.81 grease
23.83
22.80
20.62
17.89
14.47
14.22
13.20
11.93
7.53
7.39
2.54
Me
N
N
N
S
Me
Me
N
Cl
H
N
O
Et
Si
Et
O
3
250 240 230 220 210 200 190 180 170 160 150 140 130 120 110 100 90 80 70 60 50 40 30 20 10 0
f1 (ppm)
